# A Seychelles warbler genomic toolkit

**DOI:** 10.64898/2026.04.16.719046

**Authors:** Kiran G. L. Lee, Charlotte Bartleet-Cross, Sen Dong, Sergio González-Mollinedo, Alessandro Pinto, Chuen Z. Lee, Alex Sparks, Marco van der Velde, Marie-Elena Manarrelli, Tom Holden, Seychelles’ Department of Environment, Rachel Tucker, Kathryn H. Maher, Helen Hipperson, Jon Slate, Jan Komdeur, David S. Richardson, Hannah L. Dugdale, Terry Burke

**Author notes:** Corresponding Authors: Kiran Gok Lune Lee Hannah L. Dugdale Terry Burke; Terry Burke. Authors, email addresses: Charlotte Bartleet-Cross Sen Dong; Sergio González-Mollinedo; Alessandro Pinto; Chuen Zhang Lee; Alex Sparks; Marco van der Velde Marie-Elena Manarrelli Tom Holden Seychelles’ Department of Environment Rachel Tucker Kathryn H. Maher; Helen Hipperson; Jon Slate; Jan Komdeur; David Richardson; Hannah Dugdale. Joint senior authorship (Hannah L. Dugdale, Terry Burke).

## Abstract

Understanding evolutionary processes is greatly facilitated by high-quality data on genetic variation. We report the development of a genomic toolkit for a recently bottlenecked, long-term studied species, the Seychelles warbler (Ptimerl dezil; *Acrocephalus sechellensis*). This toolkit comprises a reference genome assembled into 31 chromosomes, together with functional annotations and reference-panel-free imputation of whole-genome sequences from 1,935 individuals. The genomic data have been used to assign the sequenced individuals into a genetic pedigree. Individual genomic data are associated with a suite of phenotypic metadata, amassed from three decades of fieldwork in this closed, long-term monitored population. We compared sex and parentage assigned using the genomic data with the previously recorded sex and parentage metadata to identify and correct 41 sample DNA samples labelled with the wrong identity. This population resource enables a wide range of analyses, that include, but are not limited to phylogenetics, metabarcoding, recombination rates, linkage patterns, adaptation, heritability, demographic history, selection, and inbreeding estimates. We wish to encourage interest from researchers seeking to collaborate on future analyses and data collection. Overall, our methods demonstrate the potential of next generation sequencing and statistical tools to provide dense genomic datasets at large sample sizes for wild populations.

## Introduction

Individuals, populations, species, communities and ecosystems are characterised by genetic variation. Whole-genome sequencing, which captures genetic data from across an individual’s entire genome, is one of the most powerful methods in ecology and evolution (Allendorf, 2025; Fuentes-Pardo & Ruzzante, 2017; Taylor et al., 2021; Theissinger et al., 2023). Typing millions of genetic markers throughout the genome enables a range of analyses, including on recombination rates, linkage patterns, heritability, demographic history, genetic drift, selection, adaptation and inbreeding. When paired with individual phenotypic data, whole-genome sequencing becomes a powerful tool to uncover the genetic architecture of traits (Fuentes-Pardo & Ruzzante, 2017). Monitoring natural populations over the long term enables the repeated measurement of a suite of phenotypic, behavioural and life-history traits that are being subjected to natural levels of selection and genetic drift. Such populations offer unique opportunities to turn noisy genomic data into meaningful biological interpretations. Until recently, whole-genome sequencing of hundreds of samples from “biobank”-like datasets of long-term monitored populations was prohibitively expensive. However, improvements to sequencing and bioinformatic technologies have enabled such datasets to emerge for wild populations (Cumer et al., 2021; Duntsch et al., 2021; Hasselgren et al., 2024; Stoffel et al., 2021).

Applying genomic tools in conservation is a new and rapidly developing science, known as conservation genomics (Allendorf et al., 2010). Genomic variation can be used to deduce population size, structure, demographic history, phylogeny, genetic diversity, adaptive potential and inbreeding costs (Theissinger et al., 2023). Inbreeding depression, the reduction in fitness of offspring from related parents, is expected to be particularly severe in populations that have recently shrunk or become more isolated, because their genetic load may become unmasked. Quantifying the strength of inbreeding depression and loci that contribute to this can guide the selection of candidate individuals for translocation and genetic rescue (Kardos et al., 2016). Genetic rescue should also consider the risk of outbreeding depression, the reduction in fitness from introduced masked load or alleles that are not locally adaptive (Frankham et al., 2011). The most effective conservation genomics research will translate genetic variation data into meaningful biological understanding.

Long-term intensively monitored populations therefore offers some of the best potential for such research by pairing genomic data with often rich phenotypic and environmental data that varies through time, space and across individuals (Dussex et al., 2023; Speak et al., 2024; van Oosterhout et al., 2025).

The endemic Seychelles warbler presents an excellent system in which to study evolution, ecology and conservation biology using genomic data. Over the past three decades, intensive fieldwork during biannual breeding seasons has amassed detailed life-history data for almost all birds on Cousin Island (29 hectares, 04° 19.8′ S, 55° 39.8′ E, Figure 1).

**Figure 1:**
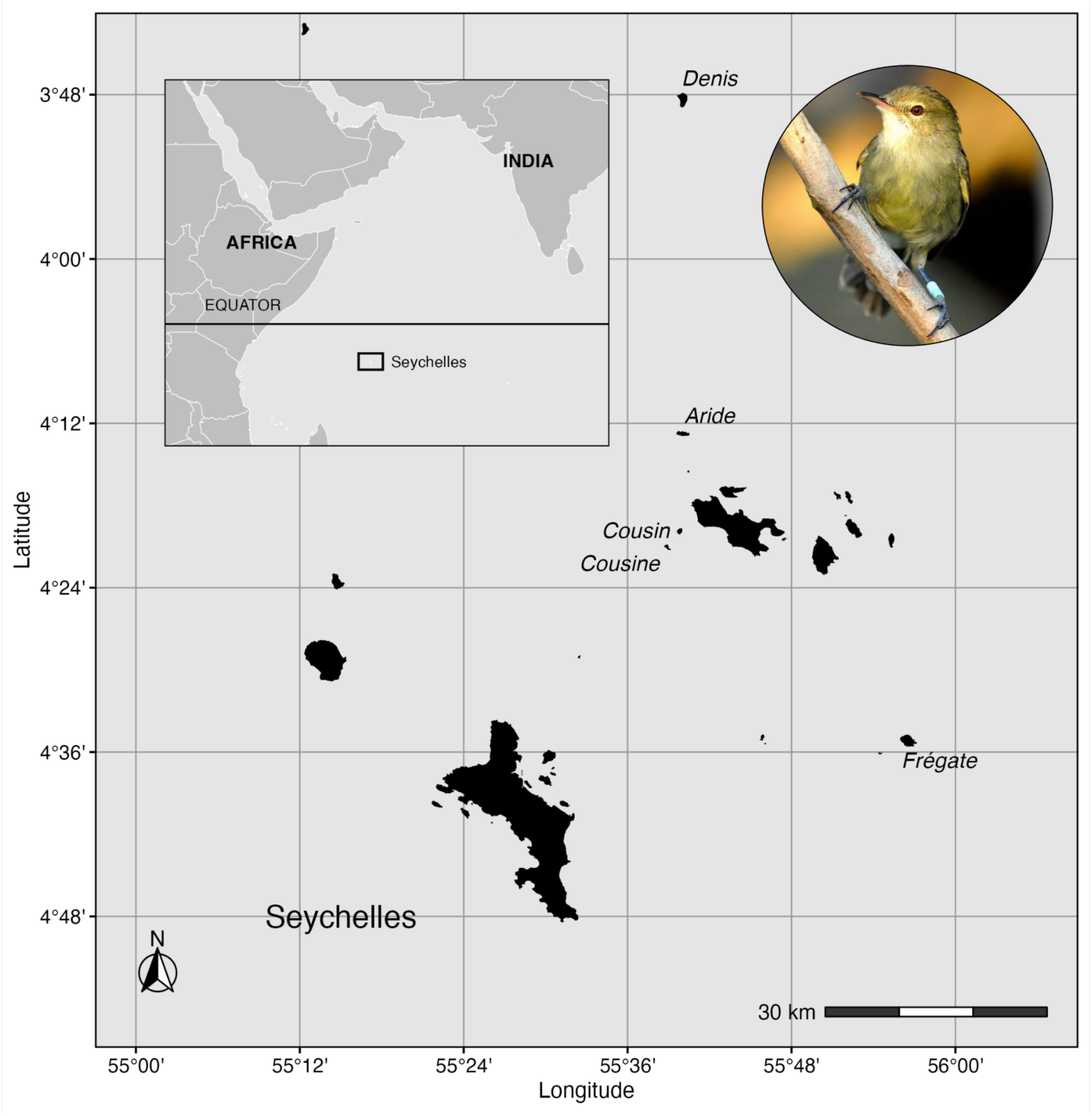
The long-term studied population of Seychelles warblers is on Cousin Island. Translocations from Cousin to Aride (1988; (Komdeur, 1994a)), Cousine (1990; (Komdeur, 1994a), Denis (2004; Richardson et al., 2006) and Frégate (2011; Wright et al., 2014) have established additional viable populations. Photograph of a colour-ringed Seychelles warbler with permission of Christopher Jean-Luc Mahoune.

Exceptionally accurate quantification of fitness traits, therefore, makes this a system well-suited to answering key questions in evolution, ecology and conservation biology. Previous research topics have included, in behavioural ecology: cooperative breeding (Komdeur, 1992, 1994b; Richardson et al., 2002; van Boheemen et al., 2019), personalities (Edwards et al., 2015, 2017), mate choice (Brouwer et al., 2010; Richardson et al., 2005; Wright et al., 2016) and sex allocation (Borger et al., 2023; Komdeur, 1996; Komdeur et al., 1997); in health: biomarker identification (Barrett et al., 2013; Brown, Spurgin, et al., 2022; Sparks et al., 2022), senescence (Berg et al., 2020; Brown, Dugdale, et al., 2022; Hammers et al., 2015), disease resistance (Hammers et al., 2016; van de Crommenacker et al., 2011; van Oers et al., 2010) and microbiome diversity (Lee et al., 2025; Worsley et al., 2022, 2025); and in conservation: impacts of translocation (Komdeur, 1994a; Komdeur & Pels, 2005; Richardson et al., 2006; D.J. Wright et al., 2014), genetic diversity (Gilroy et al., 2016; Komdeur et al., 1998; David J. Wright et al., 2014) and inbreeding (Brouwer et al., 2007; Eikenaar et al., 2008; Richardson et al., 2004). Most long-term natural study populations are based in Western Europe, North America or Australasia (Bonnet et al., 2022); by contrast, our genomic toolkit supports the study of a tropical island population susceptible to extreme variation in annual rainfall, exacerbated by El Niño events (Etongo et al., 2021; Fereday et al., 2025; Roxy et al., 2019; Tong et al., 2025).

We present a genomic dataset for Seychelles warblers that comprises a chromosome-level genome assembly, with functional annotations, and imputed low-coverage whole-genome sequences for 1,935 individuals for which we have collected a suite of associated life-history and morphological metadata from long-term monitoring. This dataset has been quality checked for integrity. The identities of archived blood and DNA samples were verified by using SNP data to assign parentage and sex, and then comparing with previous parentage and sex assignments based on microsatellite and molecular sex markers, respectively. This resource provides the opportunity for extensive downstream analyses.

Our methods use open-source software and may guide others in collecting genetic variation data in non-model systems. Our research has been done in partnership with the Seychelles’ Ministry of Agriculture, Climate Change & Environment, and long-term collaborators, Nature Seychelles (which manages Cousin Island). We recognise Seychelles’ sovereignty over the genomic data and associated metadata, and share this dataset to facilitate global collaboration in scientific research (Carroll *et al*. 2021; Jennings *et al*. 2023; McCartney *et al*. 2022; Robbins *et al*. 2023).

## Methods

### The Seychelles warbler study system

Seychelles warblers are endemic passerines that are culturally significant to the Seychelles and have been subject to intense conservation effort. Between the 1940s and 1960s, the global population of Seychelles warblers was reduced to a reported count of less than 30 within the 0.2-hectare mangroves of 29-hectare Cousin Island (04° 19.8′ S, 55° 39.8′ E, Figure 1 & Figure 2, Vesey-Fitzgerald, 1940; Crook, 1960). In 1968, BirdLife International purchased the Island, and over time replaced the coconut plantation with pan-tropical coastal vegetation, providing suitable habitat for the Seychelles warbler population to recover to 240–430 individuals (Komdeur, 1992, 1994a; Komdeur & Pels, 2005). Since 1998, a local NGO, Nature Seychelles, has managed Cousin Island. Translocations have led to viable populations on four other islands, Aride (1988; Komdeur 1994a), Cousine (1990; Komdeur 1994a), Denis (2004; Richardson et al., 2006) and Frégate (2011; Wright et al., 2014), resulting in a total, global population of >3,000 individuals (Brown et al., 2023), now listed by the IUCN only as “Vulnerable”.

**Figure 2:**
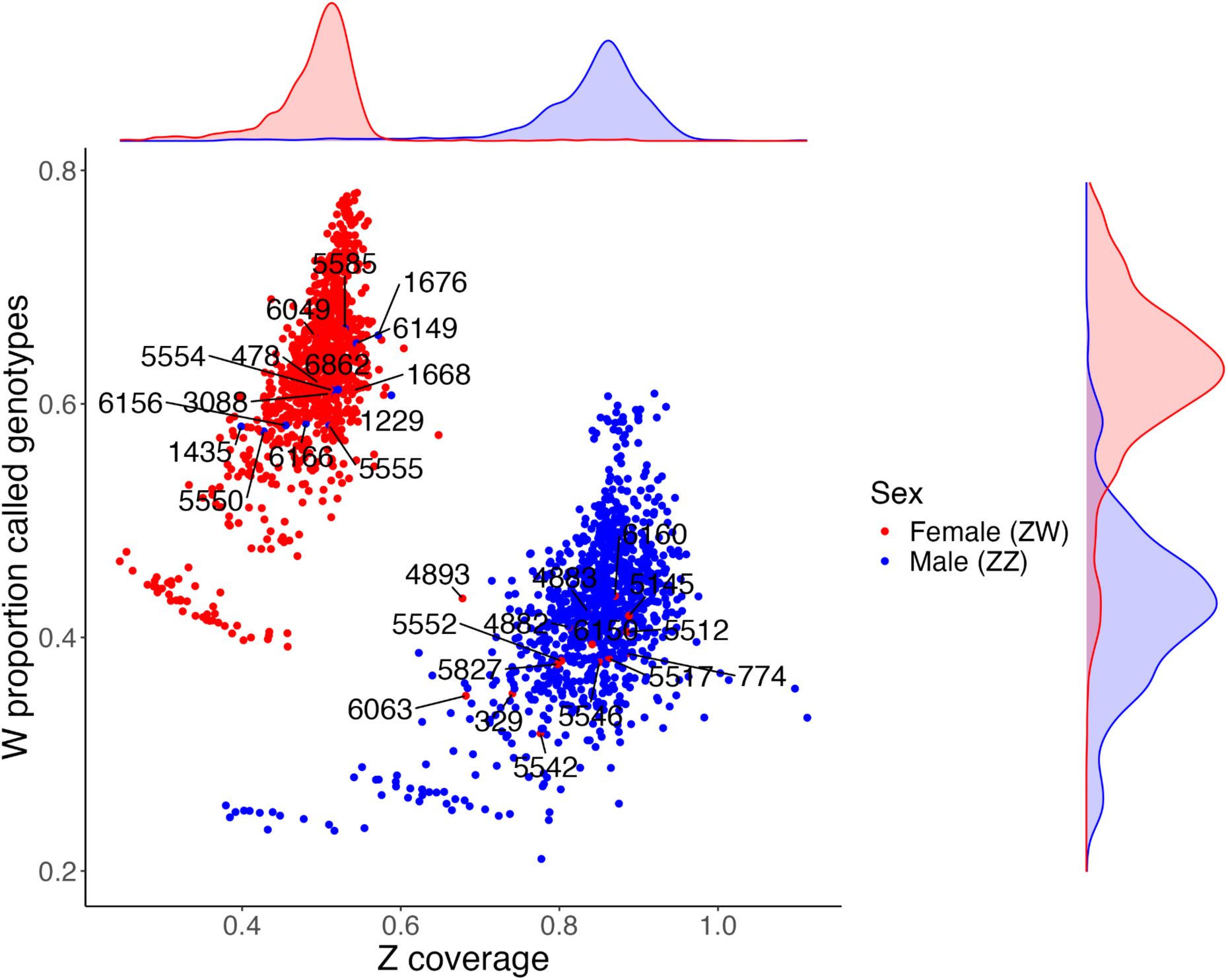
Sexing whole-genome sequenced Seychelles warblers (N = 1,976) using individual Z chromosome proportional coverage (x axis), W chromosome proportion called genotypes (y axis) and Z chromosome heterozygosity (not displayed), with 31 sex-mismatches identified and labelled according to the general linear regression model (see Sex Assignment methods).

The Cousin Island warbler population has been monitored since 1985 (Komdeur, 1992). Since 1997, >96% of all individuals have been ringed with a unique combination of a British Trust for Ornithology (BTO) metal ring and three colour rings, enabling identification of individuals (Komdeur et al., 1997; Raj Pant et al., 2020; Richardson et al., 2001). Individuals are usually first caught as nestlings, or as dependent juveniles (<5 months old) in their natal territory using mist nets (Kingma et al., 2016). These territories are often stable across time and space (Komdeur, 2001) and are defended by a breeding pair or facultatively cooperative breeding group (Komdeur, 1992, 1994b; Richardson et al., 2002; van Boheemen et al., 2019). Fieldwork takes place twice a year during the major (June to September) and minor (January to March) breeding seasons. Along with a suite of phenotypic measurements, blood samples from each capture are taken and stored in absolute ethanol at ambient temperature in the field then at −20°C in the laboratory. DNA has been extracted from blood samples each year and have been genotyped for sex determination at the *CHD* gene (Griffiths et al., 1998; Sparks et al., 2022) and for parentage assignment at 30 microsatellite loci (Edwards et al., 2017; Hadfield et al., 2006; Sparks et al., 2022; Worsley et al., 2022). More recently, faecal samples for gut microbiome research (Lee et al., 2025; Worsley et al., 2025) and blood, faecal and oral swab samples for virome analysis have also been collected. The population is small (*n* ≈ 320) and virtually contained, as only ∼ 0.10% of Seychelles warblers from Cousin have been recorded to migrate between islands in the archipelago (Komdeur et al., 2004). There are no natural adult predators making extrinsic mortality low (Komdeur & Kats, 1999). High annual resighting rates for every individual, together with accurate parentage assignment enables exceptionally accurate estimation of fitness measures such as annual survival, lifespan, annual fecundity and lifetime fecundity, making the Seychelles warbler system well-suited to answering key questions in evolutionary biology.

### Genome assembly

#### Sampling

For the reference genome we used 25 µL of blood from a female adult Seychelles warbler (BTO ring number: X784200) from Cousin Island, collected in 2017 via venipuncture and stored in 1 ml RNAlater (Ambion, USA) (Sample 1). We collected a second similar blood sample (Sample 2) from the same individual into 1 ml absolute ethanol in 2018 for scaffolding, using 10X linked reads. Additionally, to generate a population-representative consensus sequence and correct for potential individual-specific artifacts in the primary assembly, we used pooled DNA from 50 additional individuals randomly selected from the Cousin Island population. These samples represent a cross-section of the long-term monitored population, providing high-accuracy Illumina data for the final polishing stages.

#### Sample preparation

We removed salts from the RNALater sample by pelleting by centrifugation and cleaning in low TE (10 mM Tris-HCl pH 8.0, 0.1 mM EDTA). We extracted approximately 5 µg (>50 ng/µL) of high-molecular-weight DNA from each sample, using phenol-chloroform extraction based on Sambrook, Fritsch and Maniatis (2012). We sheared Sample 1 to 20 kbp using a g-TUBE (Covaris), constructed a SMRTbell library using the SMRTbell Express Template Prep Kit v2.0 (Pacific Biosciences) and selected reads greater than 15 kb using a BluePippin system (Sage Science). We constructed a Linked-Reads library for Sample 2 with no shearing or size-selection, using the Chromium Genome Sequencing Solution (10x Genomics).

We used a salt extraction protocol (Richardson et al., 2001) to extract DNA across 50 pooled samples with an average concentration of 64 ng/µL and total input of 500 ng. Samples were quantified using a Qubit 3.0 Fluorometer (Thermo Fisher Scientific, Waltham, Massachusetts, USA) with the dsDNA Broad Range Assay Kit (Thermo Fisher Scientific, Waltham, Massachusetts, USA) and DIN score of 6.8 using the Agilent TapeStation System (Agilent Technologies, Santa Clara, CA, USA). Edinburgh Genomics (University of Edinburgh, UK) constructed a library using an Illumina DNA Prep kit (Illumina, San Diego, CA, USA).

#### Sequencing

We sequenced Sample 1 on 20 SMRT cells using a Sequel Sequencing Chemistry 2.1 and Sequencing Primer v4 (Pacific Biosciences, Menlo Park, CA, USA) and Sample 2 on a NovaSeq-6000 (Illumina, San Diego, CA, USA) to produce 10X Genomics Chromium (10X, Pleasanton, CA, USA) sequenced reads. We sequenced pooled samples using 150-bp paired-end reads on 16% of an S1 lane of a NovaSeq-6000 (Illumina, San Diego, CA, USA), yielding 452 million reads.

#### Genome assembly

We assembled the reference genome using FALCON and FALCON-unzip (Chin et al., 2013), and polished it with Arrow in successive rounds, using DNANexus (Fang & Wang, 2022) to correct for errors. We used Longranger basic v2.2.2 (10X Genomics, Pleasanton, CA, USA) to convert the raw 10X Chromium reads into an interleaved file for use in ARCS v1.1.1 (Yeo et al., 2018) and LINKS v1.8.5 (Warren et al., 2015), with tigmint v.1.1.1 for error correction using the following parameters: -c 5 -m 50-10000 -s 98 -r 0.05 -e 30000 -z 500 -l 5 -a 0.3 (Jackman et al., 2018). We trimmed raw pooled Illumina data barcodes and reads for a minimum Phred quality score of 3 and a minimum length of 36 bp using Trimmomatic v0.38 (Bolger et al., 2014), and mapped 97.4% of 317,869,644 reads to the draft genome sequence with bwa v0.7.17 (Li & Durbin, 2009). We used samtools v1.11 (Li *et al*., 2009) to sort and index the sequences and polished the genome with Pilon v1.23, using the pooled reads iteratively mapped three times to the reference genome (Walker et al., 2014).

#### Evaluating the assembly

We evaluated the assembly quality after each iteration of scaffolding and polishing using BUSCO v3.1.0 (Simão et al., 2015) (Table S2). We used the vertebrate_odb9 (*n* = 2,586) gene and aves_odb9 lineage gene set (*n* = 4,915) of single copy orthologues as a benchmark for genome completeness (Simão et al., 2015). We also obtained simple assembly quality statistics using the assembly-stats package v1.0.1. We assessed non-avian contamination using blastn v2.17.0 searches to the BLAST nr nucleotide database with an e-value cutoff of 1e-25 and the -max_hsps 1 flag. We calculated read depth by mapping the 50-sample pooled Illumina reads to the complete chromosome-level assembly using bwa-mem v0.7.17, and the resulting alignments were processed with samtools v1.11. Taxonomic hits from the BLAST search and read depth data were combined and visualised in a blob plot using BlobToolKit (Challis et al., 2020).

#### Chromosome scaffolding

We scaffolded the 973 scaffolds obtained for the Seychelles warbler to the *de novo* Eurasian reed warbler (*Acrocephalus scirpaceus*) chromosome-level assembly (Sætre et al., 2021) using the scaffold function in RagTag v2.1.0 (Alonge et al., 2022; command: ragtag.py scaffold -r -C -g100 -m 100000 -f 200 -u --mm2-params ’-x asm5’). RagTag does not alter input draft assemblies but orders and orients scaffolds, connecting them with gaps guided by the known orthologous genome sequence from the other species. We visualised synteny between the Eurasian reed warbler chromosome-level assembly and the Seychelles warbler chromosome-level assembly using mcscan in the JCVI package v1.5.1 (Tang et al., 2024).

#### Genome repetitive content

We created a *de novo* repetitive DNA library for the Seychelles warbler genome using EarlGrey v4.2.4 (Command: earlGrey -g ragtag.scaffold.fasta -s Acrocephalussechellensis - o /fastdata/bop21kgl/TE/ -t 30 -d yes; (Baril et al., 2024). Earl Grey is a fully-automated transposable element (TE) annotation pipeline that also identifies repetitive sequences. Known repetitive elements from the Dfam v3.7 database (Storer et al., 2021) were identified with RepeatMasker. De novo TE identification was performed with RepeatModeler2 (Flynn *et al*., 2020). TE lengths were optimised through an automated, iterative “BLAST, Extract, Extend” process (Platt et al., 2016). This *de novo* repeat library was then annotated for TEs using RepeatMasker. Earl Grey was then used to resolve overlapping and fragmented annotations. Specific to the Earl Grey pipeline, we did not perform clustering of the TE library consensus sequences, and did not remove short TE sequences <100 bp from the annotation.

#### Gene annotation

We annotated the soft-masked genome using the fully automated gene pipeline Galba v1.0.11 (Brůna et al., 2023) with MiniProt 0.12-r237 (Li, 2023) (command: apptainer exec galba.sif galba.pl --species=’Acrocephalussechellensis’ --genome=Acrocephalussechellensis.softmasked.fasta --prot_seq=protein.faa --prg=miniprot --threads=16 --makehub --email= --crf --gff3). Galba trains AUGUSTUS (Buchfink et al., 2015; Hoff & Stanke, 2019; Stanke et al., 2006) for a novel species using miniprot and subsequently predicts genes using AUGUSTUS in the genome of that species. We used the well-characterised chicken NCBI annotation release 106 GCF_016699485.2-GdsRCg7b to build a training gene set. We used BUSCO v5.8.2 (Manni et al., 2021) aves_odb10 dataset (*n* = 8,338) to evaluate genome annotation completeness. We then performed functional annotation of the predicted proteins using eggnog-mapper v2.0.1 and the eggnog database v5.0.2 (Cantalapiedra et al., 2021; Huerta-Cepas et al., 2019) via eggnog-mapper’s web resource (command: emapper.py --cpu 20 --mp_start_method forkserver --data_dir /dev/shm/ -o out --output_dir/emapper_web_jobs/emapper_jobs/user_data/MM_d_0tlu0w --temp_dir/emapper_web_jobs/emapper_jobs/user_data/MM_d_0tlu0w --override -m diamond --dmnd_ignore_warnings -i/emapper_web_jobs/emapper_jobs/user_data/MM_d_0tlu0w/queries.fasta --evalue 0.001 --score 60 --pident 40 --query_cover 20 --subject_cover 20 --itype proteins --tax_scope auto --target_orthologs all --go_evidence non-electronic --pfam_realign none --report_orthologs --decorate_gff yes --excel; http://eggnog-mapper.embl.de/).

### Whole-genome sequenced samples

We sequenced 1,976 (972 female and 1,004 male) Seychelles warblers from the Cousin Island population at low coverage (mean = 2.65×, SD = 1.70) using blood samples collected throughout the study since 1982 (Figure S1), stored in absolute ethanol at −20°C. DNA was extracted using a salt extraction protocol (Richardson et al. 2001) or with a Qiagen DNeasy Blood and Tissue Kit. To obtain a high-coverage reference set (mean = 14.22×, SD = 2.30) we chose 56 of these individuals with no more than 2nd-degree relatedness as calculated by PLINK v2.0’s --make-king function. This set of 56 individuals was used to ground-truth the accuracy of the imputation used to obtain complete genomes of all the samples. Samples were prepared for whole-genome sequencing using the NEBNext Ultra II DNA Library Prep Kit for Illumina (New England Biolabs, Ipswich, MA), with a modified protocol designed by the NERC Environmental Omics Facility at the Centre for Genome Research (University of Liverpool) for 1/10th miniaturisation on the mosquito HV genomics platform, and were sequenced on a NovaSeq 6000 system (Illumina, San Diego, CA, USA) to generate paired-end 2 x 150-bp reads.

#### Variant discovery

We quality checked raw reads by FastQC v0.11.9 and trimmed the Illumina barcodes from reads with a minimum Phred quality score of 33 and a minimum length of 80 bp using Trimmomatic v0.39. We aligned trimmed reads to the indexed chromosome-level Seychelles warbler genome assembly using bwa-mem v0.7.17 and removed unmapped reads using samtools v1.11. We called variants with a minimum mapping quality score of 20, minimum base quality score of 20 and substitution rate of 1x10^-6^ using mpileup in bcftools v1.11 (Danecek et al., 2021), producing 18,524,883 biallelic SNPs from 1,976 samples, at mean 0.79 missingness.

#### Imputation

We imputed aligned reads from 1,976 low-coverage whole-genome sequenced samples using STITCH v1.7.0 (Davies et al., 2016) to recover missing genotypes at each of the 18,524,883 sites where biallelic SNPs were called. STITCH parameters were as follows: method = diploid, *K* = 8, and *n*Gen = 25. We conducted a sensitivity analysis to choose *K* (the number of founder haplotypes) and *n*Gen (the number of generations since founding) STITCH parameters by evaluating performance of different parameters on a single chromosome (Chromosome 10), trading off higher values of *n*Gen and *K* against exponentially increasing computational demands (Table S1). *K* and *n*Gen values were consistent with historical records (Crook, 1960; Loustau-Lalanne, 1968; Vesey-Fitzgerald, 1940) and previous demographic history analysis based on 12 microsatellite markers, that suggested a recent population bottleneck of <50 founders, 55 generations ago (Spurgin et al., 2014). We scaled recombination rates per chromosome to chromosome size (Singhal et al., 2015) following the results that Hooper *et al*. (2024) obtained using STITCH in long-tailed finches (*Poephila acuticauda*) (>25 Mbp, expRate = 1.0; <25 Mbp, expRate = 5.0, and <10 Mbp, expRate = 10.0).

We assessed the accuracy of imputation using the genotypes of the 56 reference samples. We used a conservative “test” set: the same samples downsampled to just 0.1x using samtools v1.11, followed by imputation using STITCH. We calculated genotype mean concordance between the reference and “test” sets using the bcftools gtcheck function, for each chromosome.

#### Pedigree construction

We used 572 imputed SNPs in sequoia v2.5.6 (Huisman, 2017) to assign parentage to 1,973 individuals. We selected SNPs with minor allele frequency > 0.3, genotyping rate > 99.9 and limited linkage disequilibrium (LD) (--indep-pairwise 1000 2 0.1 in PLINK v2.0; Chang *et al*. 2015), and excluded SNPs on the sex chromosomes and those on chromosomes 19 and 29, which had < 90% imputation accuracy. This filter step removed three individuals with insufficient (< 0.01×) read depth from the 1,976 imputed sequences (thus *n* = 1,973 from hereon). In sequoia, for each individual, we inputted: sex, birth date and death date, setting the maximum age at 19 years for producing offspring (Hammers et al., 2019), and assuming overlapping generations with the default genotype accuracy parameters. We assigned sex by PCR amplification of the *CHD* gene (Griffiths et al., 1998; Sparks et al., 2022). We compared parentage assignments generated using the SNP data obtained here to a previous pedigree based on microsatellite markers. This earlier “microsatellite pedigree” was created using a Bayesian Markov chain Monte Carlo (MCMC) model, MasterBayes v2.52 (Edwards et al., 2017; Hadfield et al., 2006; Sparks et al., 2022; Worsley et al., 2022) using genotypes from up to 30 microsatellite markers (Richardson et al., 2001) and key life-history data – natal territory, breeding status, age and territory distance – to assign relationships which were then filtered for posterior probability > 80% (Edwards et al., 2017). To evaluate improvements made by the sequoia pedigree, we used pairwise genomic relatedness of sequenced individuals and plotted these against pairwise relatedness estimates from both the microsatellite-based pedigree and sequoia pedigree. Pairwise genomic relatedness was calculated using GCTA (Yang et al., 2011) and pairwise pedigree-based relatedness was calculated in the R package kinship2 (Sinnwell et al., 2014).

#### Sex assignment

We assigned sex to whole-genome sequences (*N* = 1,973) using (1) the proportion of called W chromosome genotypes (where males are expected to have a lower value), calculated using PLINK v2.0’s function --missing, (2) Z chromosome coverage scaled to genome-wide coverage where males (two copies of the Z) are expected to have a higher value than females (one copy of the Z), calculated using the idxstats function in samtools v1.11 and dividing by mean sample coverage, and (3) Z chromosome heterozygosity (where females are expected to have values close to zero), calculated using PLINK v2.0’s function: --sample-counts cols=hom,het flag. We performed a logistic regression, using the glm function in the R package, stats v4.4.3, of how previously assigned sex by *CHD*-gene typing, could be predicted by variation in (1), (2), the interaction between (1) and (2) and (3), assuming a binomial distribution (Table S4). We validated the model’s ability to discriminate between males and females using the Area Under the Receiver Operating Characteristic curve (AUC), demonstrating “excellent” performance with an AUC of 0.989 (Çorbacıoğlu & Aksel, 2023). We checked for conflicts in sex prediction by *CHD* gene and by SNP data by visually inspecting the plot of (1) against (2), coloured for *CHD*-gene sex (Figure 2).

#### Sample audit

We followed the framework described by Duntsch *et al*. (2022), to verify the identities of our samples, using sex and pedigree meta-data to identify and correct sample mismatches.

Out of 1,976 individuals we found inconsistencies in parentage (*n* = 100, 5.2%, with at least one mismatched parent) and sex (*n* = 31, 1.6%) to their previous assignments. We also found unintentional duplicates (*n* = 29, 1.5%) using PLINK v2.0’s flag --king-table-filter to estimate KING kinship coefficients among all samples (Manichaikul et al., 2010), verified in sequoia v2.5.6 (Huisman, 2017). When looking for commonalities of these mismatched samples with their phenotypic data, 15/29 duplicate pairs, then 12/20 de-duplicated sex-mismatched samples, and 14/151 de-duplicated, sex matched but parentage-mismatched samples had been obtained from a single plate of 50 samples whose DNA had been extracted in an earlier project (Plate “X”). We describe the process for correcting sample mismatches in the supplementary methods as summarised in Figure 3. All sample decisions were verified by a second reviewer and decision conflicts were resolved by a third reviewer (Tables S7, S8).

**Figure 3.**
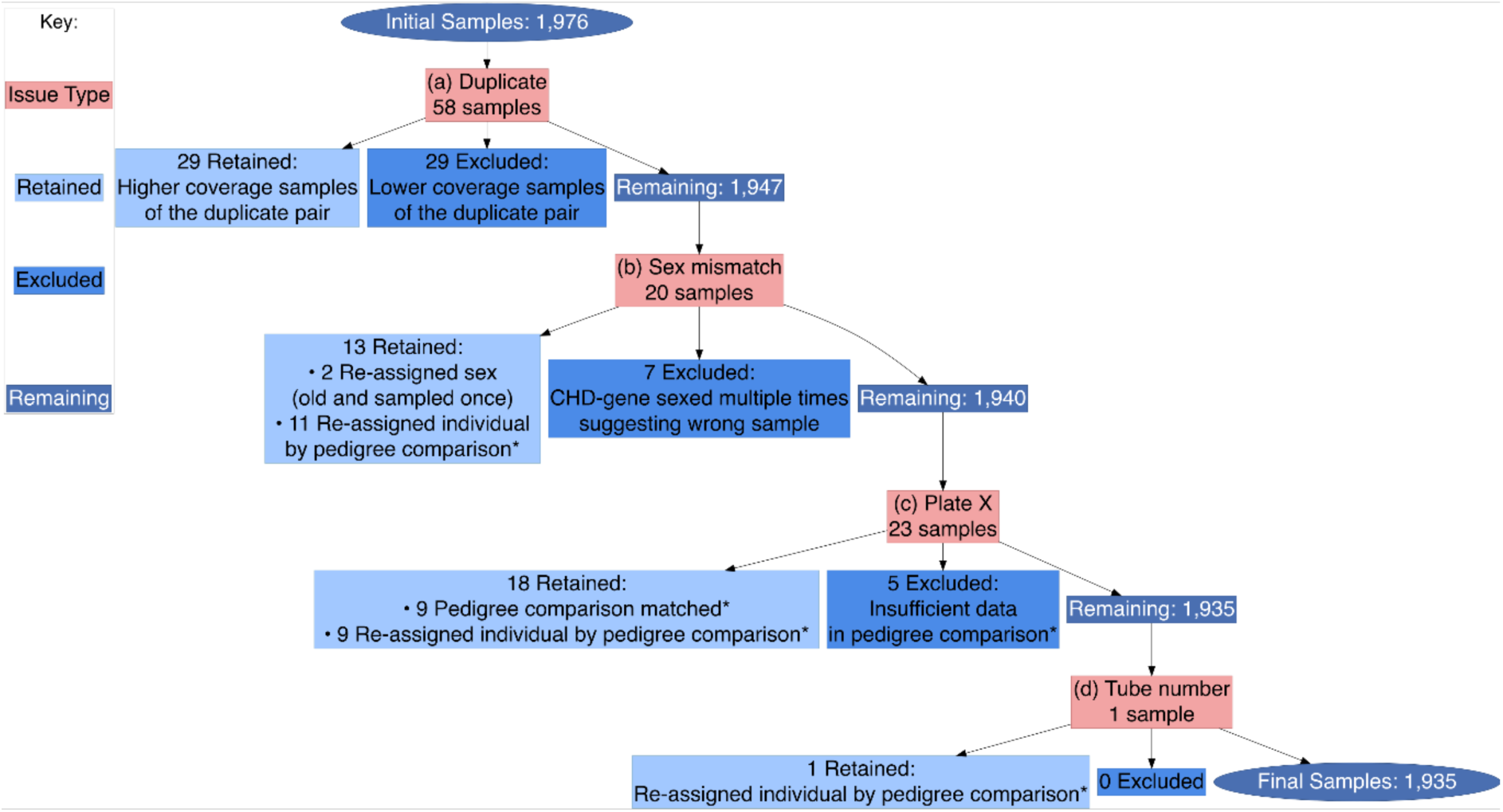
Flow diagram detailing action taken for identified mix-ups to leave 1,935 Seychelles warbler individual samples with verified identities and whole-genome sequence data. * Pedigree comparison involved comparison of parent assignment by sequoia using SNP data with previous assignment by Bayesian analysis of microsatellite genotypes and meta-data.

### Mitogenome

We assembled a mitochondrial genome using a hybrid assembly approach, leveraging structural information from PacBio long reads and incorporating the accuracy of Illumina short-reads.

#### Read filtering

Illumina short reads: 150-bp paired-end Illumina reads from 56 reference individual samples were pooled *in silico*. To isolate mitochondrial reads from this meta-pool, we first mapped all reads to the comprehensive animal_mt database from GetOrganelle v1.7.7.1 using bowtie2 v2.3.5. Read pairs where both mates were successfully mapped were retained using samtools v1.11. We synchronised the resulting FASTQ files using the repair.sh script from the BBMap suite v38.90.

PacBio Long Reads: Raw PacBio subreads from a single individual (Sample 1) were aligned to the mitochondrial genome of a congeneric species, the Oriental reed warbler (*Acrocephalus orientalis*, NC_059273.1), using pbalign v0.4.1 and extracted into a clean FASTQ file using samtools v1.11.

#### Assembly and Finishing

Filtered mitochondrial Illumina short reads and PacBio long reads were used as input for a hybrid assembly with Unicycler v0.5.1 in --mode bold. This produced a single, complete, circular contig of 17,847 bp.

#### Annotation and Validation

The final circular contig was annotated using MitoZ v2.4 and visualised as a Circos plot (Krzywinski et al., 2009). The annotation identified all 37 expected genes: 13 protein-coding genes, 22 transfer RNA (tRNA) genes, and 2 ribosomal RNA (rRNA) genes. To validate the assembly, the gene order was compared to the Oriental reed warbler reference genome (Kim et al., 2020).

## Results

### Genome assembly

Our Seychelles warbler genome assembly totalled 1.081 Gbp in length. Assembly properties are described in Table 1. In total, 1.072 Gbp (98.25% of the genome) were assigned to 29 autosomes and 2 sex chromosomes using RagTag (Figure 4a, Table S3). Of this, 0.019 Gb (1.75% of the genome) could not be assigned. Chromosome 19 was poorly assembled, comprising 206,394 base pairs. Visualisation of synteny plots between the great tit (*Parus major*) and both the Seychelles warbler and reed warbler showed homology between chromosome 19 of the reed warbler and chromosome 4a of the great tit (Figure 4b), whereas chromosome 17 of the Seychelles warbler showed homology to 4a of the great tit (Figure 4c). We did not detect non-avian contaminant sequences in the assembly using blobtools (Figure S2)

**Figure 4.**
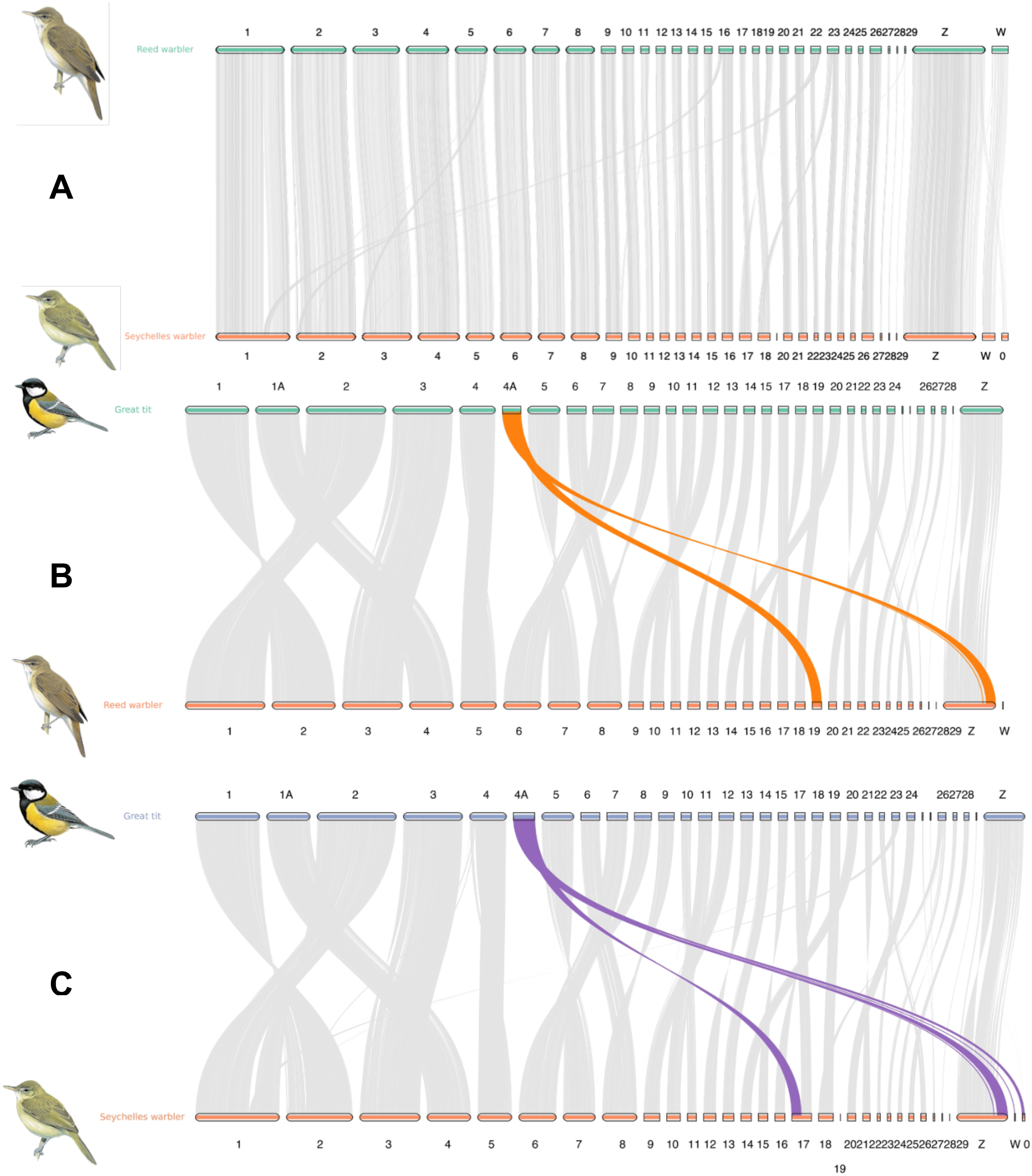
Chromosome assembly synteny plots between (A) reed warbler (Acrocephalus scirpaceus) and Seychelles warbler (Acrocephalus sechellensis), (B) great tit (Parus major) and reed warbler and (C) great tit and Seychelles warbler.

**Table 1.** Basic genome assembly statistics for the Seychelles warbler.

| Metric | Value |
| --- | --- |
| Number of scaffolds | 973 |
| Total size of scaffolds (bp) | 1,081,018,985 |
| Useful ($\geq 25$ kbp) amount of scaffold sequences (bp) | 1,078,983,347 |
| Longest scaffold (bp) | 59,560,152 |
| Number of scaffolds $> 1$ kbp | 910 |
| Number of scaffolds $> 10$ kbp | 874 |
| Number of scaffolds $> 100$ kbp | 461 |
| Number of scaffolds $> 1$ Mbp | 110 |
| Number of scaffolds $> 10$ Mbp | 29 |
| N50 (bp) | 18,737,313 |
| L50 | 16 |
| NG50 (bp) | 21,515,182 |
| LG50 | 14 |
| %A | 28.9 |
| %C | 21.1 |
| %G | 21.1 |
| %T | 28.9 |
| Total Number of Ns | 36,729 |
| %N | 0.003 |
| BUSCO (vertebrate lineage) | 96.8% |
| BUSCO (avian lineage) | 93.3% |

#### Genome repetitive content

Annotation of repeated sequences using EarlGrey identified that 11.5% of the female Seychelles warbler genome assembly consisted of repetitive sequences (Figure S3). Analysis of element subfamilies revealed that long interspersed nuclear elements (LINEs), specifically CR1, and long terminal repeats (LTRs), including ERVL, ERV1, and ERVK, were the most abundant (Figure S3).

#### Gene annotation

We predicted 20,080 protein-coding genes across the 29 autosomes and 2 sex chromosomes, with 96.4% genome annotation completeness as evaluated by BUSCO (Table S5).

#### Mitogenome

The complete mitochondrial genome of the Seychelles Warbler was assembled into a single, circular contig of 17,847 bp (Figure S4). The overall base composition was A: 29.3%, C: 32.3%, G: 15.4%, T: 23.0%, with a total GC content of 47.7%. The sequence exhibited a positive AT-skew (0.121) and a negative GC-skew (-0.355), as is characteristic of avian mitogenomes (Saccone et al., 1999). Gene order showed 100% synteny with the Oriental reed warbler, the closest relative with a complete and annotated mitochondrial genome available.

### Whole-genome sequenced samples

#### Imputation

We called variants in 1,976 samples with mean coverage of 2.6× and imputed missing genotypes using a reference panel-free approach using STITCH. This generated 18 million biallelic, genome-wide SNPs, with mean SNP missingness of 0.055 and accuracy of 96.2%, assuming 0.1× initial coverage. Imputation accuracy decreased with chromosome size (Figure S5, Table S6).

#### Sample audit

We used previous sex determination by PCR amplification of the *CHD* gene combined with parentage assignment from a previous pedigree assembled using microsatellites and Bayesian analysis of field variables to correct the identities of 40 samples (29 duplicate samples which were excluded, 11 sex-mismatched samples and 9 sex-matched but parentage mismatched samples), and we corrected 2 incorrectly sexed samples. We excluded eleven samples completely as there was no clear consensus of the correct sample identity. Of the duplicate and sex-mismatched DNA samples, 66.0% (31/47) were from a microplate of 50 samples prepared within a separate piece of research over a decade ago; a mix-up likely occurred during aliquoting of samples into this plate.

After identifying and re-assigning or removing sample mismatches, 1,935 (98.0%) whole-genome sequenced Seychelles warbler samples remained in the “cleaned dataset”, suitable for downstream analyses in future studies. We re-ran this cleaned dataset a final time for parentage assignment using sequoia to create a corrected pedigree. This sequoia pedigree showed a 98.4% assignment match rate (1,507/1,523 mothers, 1,558/1,574 fathers; 30 focal individuals had at least one parent mismatching and 0 had both mismatching) with the existing microsatellite-based pedigree.

#### Pedigree assignment

After correcting or removing mismatched samples, we constructed a *de novo*, SNP pedigree in sequoia that assigned 1,935 individuals to 1,661 dams and 1,677 sires. We assigned both parents to 1,540 individuals (mean birth date: December 2007, SD: 2,835 days), one parent to 258 individuals (mean birth date: August 1994, SD: 2,318 days) and no parents to 134 individuals (mean birth date: January 1992, SD: 1,922 days). Naturally, ancestral individuals could not be assigned parents. Correlations of genomic pairwise relatedness and pedigree relatedness between samples showed that we improved on the previous microsatellite pedigree (Figure 5a) by using SNP data in the sequoia pedigree (Figure 5b; Pearson’s r increased from 0.55 to 0.57).

**Figure 5:**
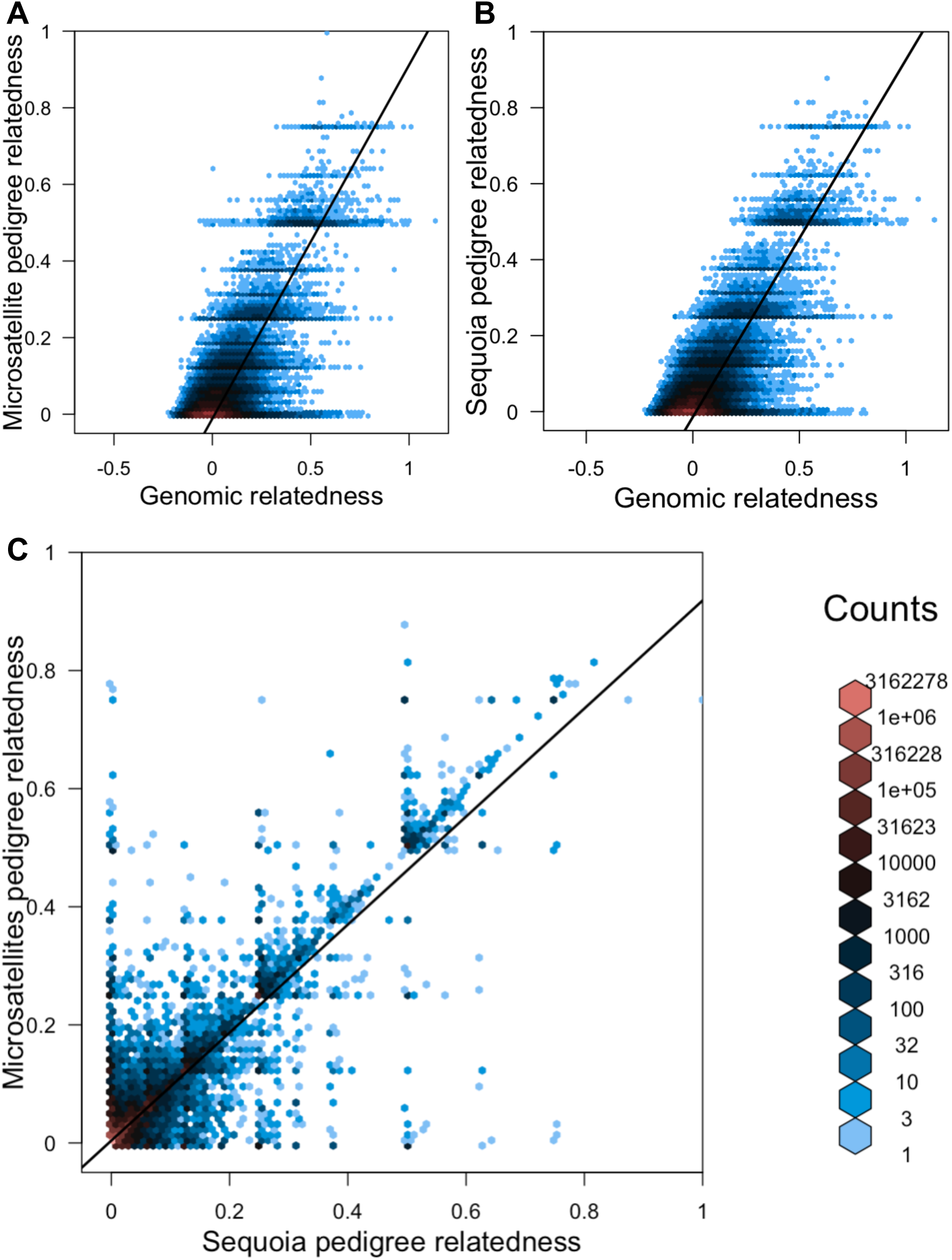
Pairwise hexbin plots A. genomic relatedness against microsatellite-based pedigree relatedness (Pearson’s r = 0.55), B. genomic relatedness against sequoia pedigree relatedness (Pearson’s r = 0.57) and C. sequoia pedigree relatedness against microsatellite-based pedigree relatedness (Pearson’s r = 0.87) in 1,932 identity-verified Seychelles warblers.

## Discussion

### Genome assembly

The Seychelles warbler genome assembly is typical in size (1.07 Gbp), contig N50 (18.7 Mbp) and BUSCO score (96.8% vertebrate lineage, 93.3%, avian lineage) relative to other avian genome assemblies (Feng et al., 2020; Rhie et al., 2021; Zhang et al., 2014). The slightly lower completeness score for the avian lineage likely reflects the larger gene set in the avian database (4,915 vs 2,586 genes) and the many avian-specific conserved genes localised to microchromosomes that are repeat-rich and difficult to assemble (Peona et al., 2021). We used the Eurasian reed warbler *de novo* chromosome-level genome (Sætre *et al*., 2021) to assemble 973 scaffolds of the Seychelles warbler genome into 29 autosomes and 2 sex chromosomes. We justified this, as karyotypes are highly conserved among birds, relative to other vertebrates (Waters et al., 2021) and the Eurasian reed warbler is congeneric with the Seychelles warbler. Chromosome 19, the tenth smallest chromosome in the Eurasian reed warbler, was poorly assembled. In the Eurasian reed warbler, chromosome 19 shows close synteny with a portion of chromosome 4A of the great tit, *Parus major* (Figure 4b). The remainder of chromosome 4A shows close synteny with the Z chromosome in the Eurasian reed warbler and is common to all studied Sylvioidea (Ponnikas et al., 2022). The dynamic nature of chromosome 19, or chromosome 4A, as it was before the most recent common ancestor to *Sylvioidea* (Ponnikas et al., 2022), may make it difficult to assemble using homology alone. As the Seychelles warbler assembly relies on the *de novo* Eurasian reed warbler assembly, scope for comparative genomic analyses may be limited. We recommend future work to produce a more complete *de novo* chromosome assembly for the Seychelles warbler using linkage, physical, or spatial proximity maps (from Chromatin Conformation Capture). Downstream analyses using this genomic toolkit can exclude chromosome 19, if necessary.

In addition to the nuclear genome, we assembled the complete mitochondrial genome of the Seychelles warbler. Typical of passerines (Kessler & Avise, 1985), the mitogenome is approximately 17.8 kbp in length and contains the standard suite of 13 protein-coding genes, 22 tRNAs, and two rRNAs. This facilitates future research into mitochondrial-nuclear co-adaptation and maternal inheritance patterns in this long-term monitored population and for future phylogenetic studies.

### Repeat content and functional annotation

A repeat content of 11.5% is typical of birds (mean = 9.5%, SD = 3.1, *N* = 363 species, Feng et al., 2020), but is likely to be an underestimate, as repetitive regions are more difficult to assemble (Benham et al., 2024). In the soft-masked genome (excluding repetitive sequences), 20,008 predicted protein-coding genes is typical of bird genomes annotated using the GALBA pipeline (Bailey et al., 2023; Chrysostomakis et al., 2025; Pawar et al., 2025) and will help the interpretation of the impact of imputed variants produced in this dataset in downstream analyses. This could be useful, for example, in linking variants to host expression pathways in understanding blood-based disease pathogens or immunity phenotypes in the Seychelles warbler (Davies et al., 2021; Hammers et al., 2016; Worsley et al., 2022).

### Imputation

We imputed 18 million biallelic variants from 1,935 individual Seychelles warblers sequenced at low (mean 2.6×) coverage. Imputation of low-coverage whole-genome sequenced samples increasingly recognised as a cost-effective alternative for generating high-density genotypes in wild species (Fuller et al., 2020; Gao et al., 2020; Stoffel et al., 2021; Wall et al., 2016; Yang et al., 2024). While most imputation frameworks require a high-coverage reference panel to identify ancestral haplotypes (Watowich et al., 2023; Wragg et al., 2024), such panels are often logistically unattainable or fail to capture the specific standing variation of natural populations. To circumvent this, we used the reference-panel-free approach in STITCH (Davies et al., 2016) which can outperform even reference-panel imputation softwares deducing haplotypes in small, inbred populations (Vi et al., 2025). The Seychelles warbler is particularly well-suited to this approach; as a closed population with a severe historical bottleneck, it possesses limited haplotype diversity per locus (Spurgin et al., 2014), and high LD (*r*^2^ = 77 kbp) with a large number of sequenced samples. These population characteristics likely facilitated a validation accuracy of 96.1% at a drastically downsampled coverage of 0.1×. Given that our study’s actual mean coverage (2.6×) was over 25 times higher than this test threshold, our final genotype calls likely exceed this accuracy level. We did observe that smaller chromosomes were imputed with slightly less precision, a likely consequence of higher recombination rates causing LD to decay more rapidly over physical distance, but the overall high resolution of this dataset provides a robust foundation for the downstream population and functional genomic analyses.

### Sample audit

By cross-referencing recorded meta-data with whole-genome data, we were able to perform a genomic audit, providing a level of certainty in individual identity to our samples. Sample mix-ups are common and somewhat unavoidable, especially for long-term field studies (Broman et al., 2015; Duntsch et al., 2022; Lippi et al., 2017; Lobo et al., 2021; Lynch et al., 2012), but are difficult to detect and underacknowledged. The concentration of errors (66.0%) within a single decade-old microplate highlights a vulnerability potentially during the aliquoting phase of transferring samples into high-density formats. Using genomics, errors which may have been difficult to detect in smaller datasets can be identified. The remaining total of 16 out of 1,926 (< 1%) samples that were found to be duplicates or mismatching the recorded sex suggests that the dataset was otherwise extremely accurate. This testifies to the overall accuracy of field and laboratory protocols maintained over the 37-year duration of sampling within the Seychelles Warbler Project. We encourage researchers to incorporate similar systematic checks, such as our comparison of heterozygosity-based sexing with field records and SNP-based relatedness with pedigrees, in their own systems and for this, recommend the guide outlined by Duntsch *et al*. 2022.

### Pedigree

An initial round of parentage assignment by sequoia showed low disagreement (5.2%, 100 individuals with one mismatching parent) with the earlier microsatellite-based pedigree before mixups were corrected or excluded. This high degree of consensus justified the use of the previous microsatellite pedigree-assigned parents to verify the correct identities of mislabelled samples in the genomic sequencing dataset. In the cleaned dataset, agreement with the microsatellite pedigree increased to 99.0% (31 individuals had one mismatching parent). This indicates that the pedigree data used in previous Seychelles warbler research was accurate (Borger et al., 2023; Brown, Dugdale, et al., 2022; Chesterton et al., 2024; Groenewoud et al., 2018; Raj Pant et al., 2019, 2020; Sparks et al., 2022). The genomic pedigree, making use of hundreds more markers, was even more accurate, as the Bayesian framework in the previous microsatellite-based pedigree incorporated data on territory locations of the parents and offspring, parents’ ages and the father’s social status that must be subject to some error, and microsatellite genotyping is also error-prone (Dakin & Avise, 2004). Any dispersal before first capture will have limited the accuracy of natal territorial assignments – as was the case for a fraction (23/31) of the previously assigned individuals that had parents mismatching in the sequoia analysis. Our future research on Seychelles warblers can now incorporate the verified genomic pedigree in new analyses on topics including genotype phasing, effective population size, inbreeding, genomic prediction and lifetime reproductive success.

### Sexing

We can use whole-genome sequence data to sex individuals (Bilton et al., 2019; Denoyelle et al., 2021; Duntsch et al., 2022). In monomorphic, heterogametic species, sex is typically assigned by PCR-amplification of orthologous genes found on each sex chromosome, dimorphic in length and distinguishable by gel electrophoresis. These markers can be prone to errors such as drop out (Pompanon et al., 2005). Whole-genome sequencing data enables sex and parentage to be assigned, and offers more scope for downstream analyses, skipping the need for separate specific assays. A logistic model using the Z chromosome coverage, the number of genotypes assigned to the W chromosome, and Z chromosome heterozygosity in each individual identified sample mix-ups, and after correcting for these, we found no sex-mismatches in the cleaned Seychelles warbler dataset. This method could be applied in other heterogametic systems by aligning reads to the most closely related species with an available chromosome-level assembly.

## Conclusions

In conclusion, this study establishes a high-quality, chromosome-level genomic toolkit for the Seychelles warbler, providing a robust foundation for future evolutionary and conservation research. The implementation of a reference-panel-free imputation strategy via STITCH proved exceptionally successful, yielding 18 million biallelic variants mapped using 29 chromosomes with contig N50 of 18.7 Mbp, and across 1,935 individuals with a minimum accuracy of 96.1% as determined from 0.1× coverage sites. This approach demonstrates that low-coverage whole-genome sequencing (mean 2.6× in our samples) is a cost-effective and highly viable strategy for the long-term monitoring of a closed, wild population. This transition to a genomic approach has provoked an essential and valuable audit of the long-term dataset. By identifying and correcting sample mismatches and sexing errors, many of which were traced to a single historical plate, we have reduced the error rate to <1%, resulting in an extremely accurate dataset. The 99% agreement between our new genomic pedigree and the historical microsatellite-based pedigree not only validates decades of previous research, but also enhances our ability to perform high-resolution analyses. We demonstrate how an integrated genomic approach can bypass the need for separate molecular assays by simultaneously capturing essential metadata, such as sex and parentage, from a single sequencing effort. This toolkit can enable analyses including genotype phasing, effective population size trajectories, and the precise genomic architecture of inbreeding depression, ultimately supporting the evidence-based conservation of this iconic island endemic. Beyond the Seychelles warbler, our methods offer broad-scale applicability for the study of non-model organisms in the wild.

## Supplementary methods

### Sample audit

#### Duplicates

Assuming that one of the two labels for each identically labelled pair was correct, we confirmed the identities of 11/29 because the sexes in the candidate pair differed, and one individual was of the correct sex. We confirmed the other 18/29 samples through concordance between the previous microsatellite and current genomic pedigree assignments to its mother (15/29) or offspring (3/29) (Figure 3a). We kept the higher-coverage sequence of the two samples in a confirmed duplicate pair.

#### Sex-mismatches

Of the 20 remaining non-duplicated sex-mismatched sequences, we assigned the correct individual identity to 11/20 sequences through concordance between the previous microsatellite and current genomic parentage assignments. We excluded 7/20 sex-mismatched samples that had been *CHD*-gene sex typed multiple times and corrected the sex for 2/20 samples that had been *CHD*-gene sexed only once and assumed to have been previously sexed incorrectly (Figure 3b).

#### Plate X mismatches

As 66.0% (31/47) of duplicate and sex-mismatched samples could be traced to Plate X, we undertook an audit of the remaining 14 de-duplicated but sex-matched samples on Plate X by comparing their assigned parents in the microsatellite and sequoia pedigrees. We corrected 9/14 samples because the genomic pedigree-assigned parents matched the microsatellite pedigree-assigned parents of the likely correct individual. We excluded the remaining 5/14 samples that had insufficient pedigree data to correctly re-assign their sample identity (Figure 3c).

#### Parentage mismatches

After correcting these mismatches, we constructed another pedigree in sequoia. To evaluate performance, we compared this with the microsatellite pedigree made using MasterBayes. This identified 35 focal samples that had at least one inconsistent, formerly assigned parent. To identify which of the assigned parents was correct, we compared key metadata: natal and parental territories, natal and parental breeding groups, and genomic relatedness (as calculated from PLINKv2.0’s --make-grm function) to the individual. The sequoia-assigned parent of 26/35 samples held more compatible metadata (same territory, or same breed group ID, or better genomic relatedness), but in 5/35 samples there was not sufficient metadata to identify the correct parent. The same pair of mismatched and correct mothers were associated with 4/35 offspring samples. These two potential mothers had the same blood tube sample identifier, a unique number assigned to both blood samples, and had each been sampled only once. This suggested a sample labelling mix up in one direction, and so we reassigned the correct mother using the sequoia assignment (Figure 3d).

## Supplementary Tables

**Table S1.** Sensitivity analysis when choosing STITCH parameters K (the number of founder haplotypes) and nGen (the number of generations since founding). r^2^ is a metric of imputation accuracy generated by STITCH following each imputation run.

| $r^2$ | $K = 4$ | $K = 10$ | $K = 30$ |
| --- | --- | --- | --- |
| nGen= 20 | 0.972 | 0.976 | 0.977 |
| nGen= 50 | 0.973 | 0.976 | 0.977 |

**Table S2.** Assembly metrics across stages of the assembly process. The size of the assembly, number of contigs, N50, number of gaps and the number of ‘Ns’ introduced into the assembly by gaps. BUSCO score (V) is the percentage of complete BUSCOs using the vertebrate lineage (n = 2,586), with BUSCO score (A) based on the Aves lineage (n = 4,915).

| Stage | Size (bp) | No. of contigs | N50 | Gaps | N count | BUSCO score (V) | BUSCO score (A) |
| --- | --- | --- | --- | --- | --- | --- | --- |
| Falcon | 1,081,153,341 | 1,404 | 4,651,707 | 0 | 0 | 92.9 | 88.1 |
| ARCS 1 | 1,081,202,141 | 1,050 | 12,516,622 | 488 | 48,800 | 92.9 | 88.1 |
| ARCS 2 | 1,081,209,042 | 990 | 15,207,662 | 558 | 55,701 | 92.9 | 88.2 |
| ARCS 3 | 1,081,211,746 | 973 | 18,737,720 | 589 | 58,405 | 93.1 | 88.1 |
| Pilon 1 | 1,081,119,075 | 973 | 18,737,359 | 566 | 41,541 | 96.8 | 93 |
| Pilon 2 | 1,081,003,470 | 973 | 18,736,971 | 559 | 38,733 | 96.8 | 93.2 |
| Pilon 3 | 1,081,010,066 | 973 | 18,736,959 | 558 | 37,443 | 96.8 | 93.3 |

**Table S3.** Chromosome sizes of the Seychelles warbler genome assembly

| Chromosome | Size (bp) |
| --- | --- |
| 1 | 159731717 |
| 2 | 122937904 |
| 3 | 120275107 |
| Z | 91483344 |
| 4 | 75128064 |
| 5 | 66803114 |
| 6 | 71256953 |
| 7 | 63941626 |
| 8 | 63476739 |
| 9 | 22849264 |
| 10 | 21752670 |
| 11 | 20740192 |
| 12 | 18966161 |
| 13 | 17000646 |
| 14 | 15744861 |
| 15 | 14404183 |
| 16 | 12042549 |
| 17 | 21860121 |
| 18 | 16211747 |
| 19 | 206394 |
| 20 | 9306870 |
| 21 | 8335059 |
| W | 9299204 |
| 22 | 2447251 |
| 23 | 3111230 |
| 24 | 6129322 |
| 25 | 6158279 |
| 26 | 5422578 |
| 27 | 2347831 |
| 28 | 2162601 |
| 29 | 511510 |
| Unassigned | 19139384 |

**Table S4.** Logistic regression results for the genomic sex determination model.

| Predictor | Estimate ($\beta$) | Std. Error | z-value | p-value |
| --- | --- | --- | --- | --- |
| Intercept | 12.38 | 3.87 | 3.2 | 0.001 |
| (1) W Proportion Non-Missing | -40.67 | 10.21 | -3.99 | < 0.001 |
| (2) Proportional Z Coverage | -1656 | 1003 | -1.65 | 0.099 |
| (3) Z Heterozygosity | -4.33E-05 | 2.25E-05 | -1.93 | 0.054 |
| (1)*(2) Interaction | 8857 | 2399 | 3.69 | < 0.001 |

**Table S5.**
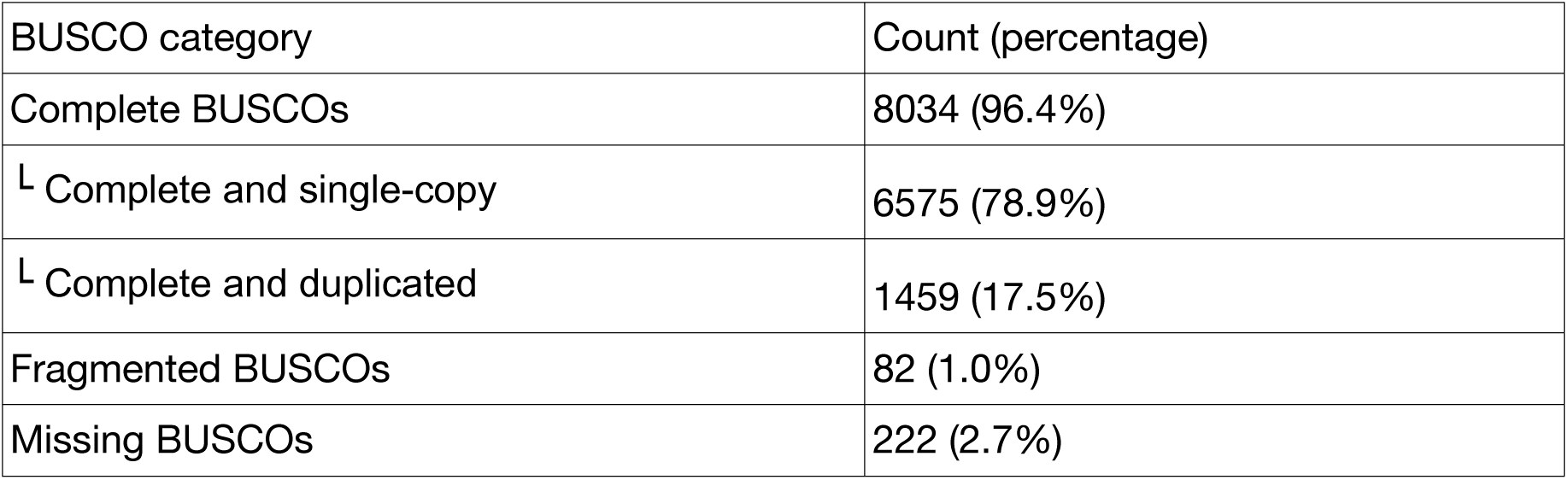
Completeness of the predicted proteome (BUSCO v5.8.2) using the aves_odb10 (n = 8,338 orthologues) dataset.

**Table S6.**
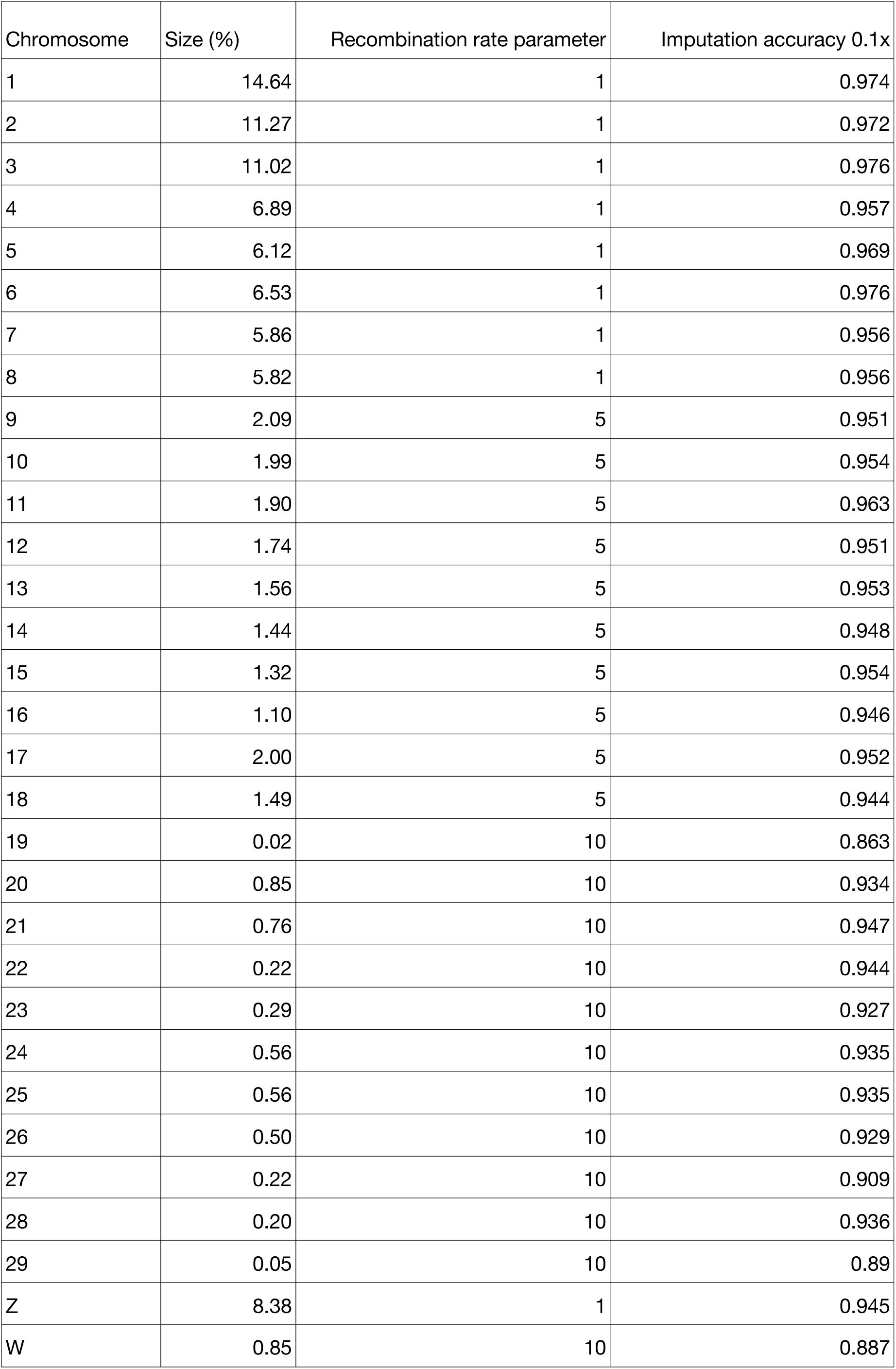
Accuracy of imputation by STITCH per chromosome.

**Table S7.**
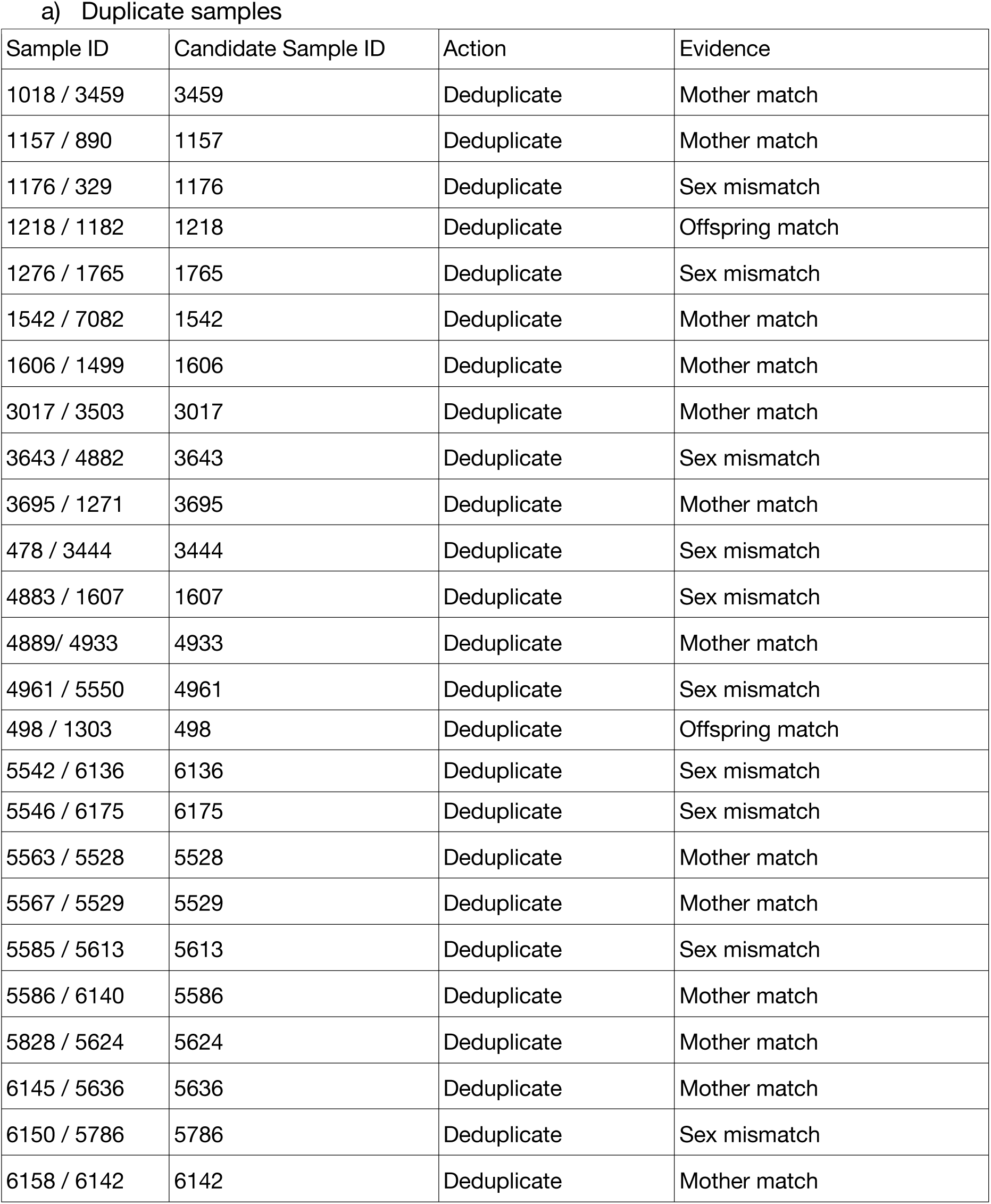
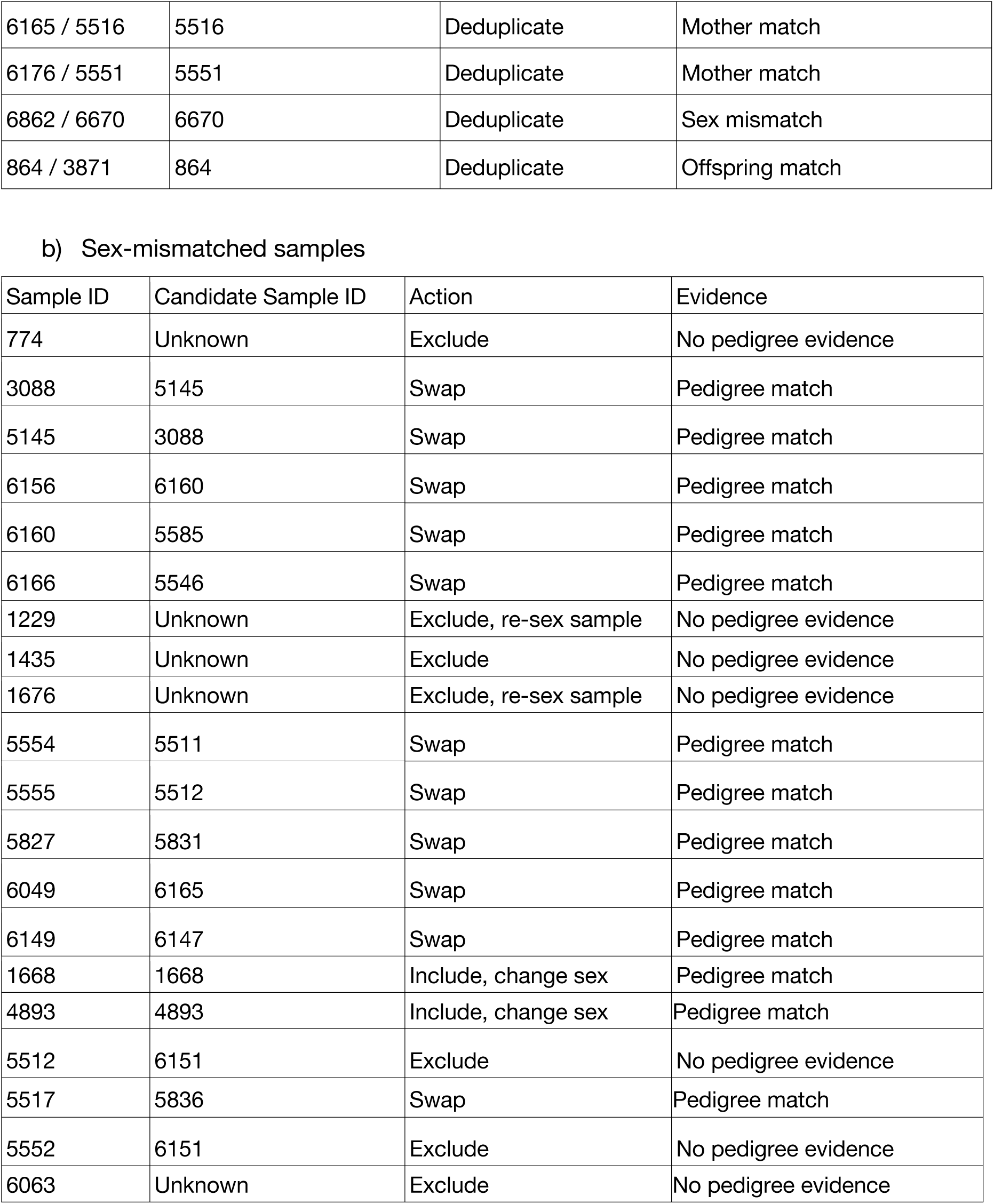
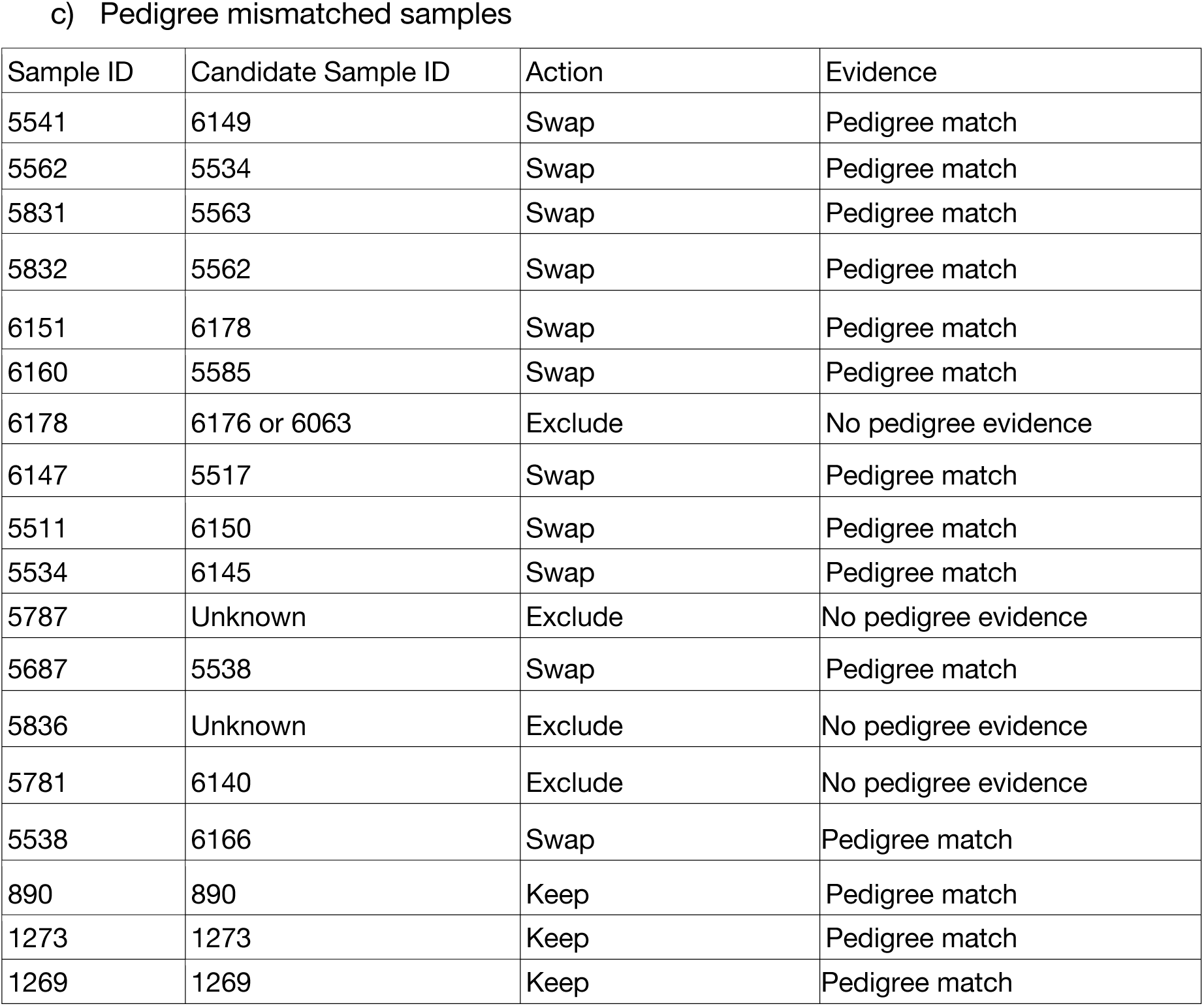
Resolution of sample identity conflicts for: a) duplicate, b) sex mismatched (non-duplicate) and c) pedigree mismatched (non-sex mismatched) samples. Decisions were based on the consensus between three independent reviewers using genomic, behavioural, and pedigree data.

| Sample ID | Candidate Sample ID | Action | Evidence |
| --- | --- | --- | --- |
| 1018 / 3459 | 3459 | Deduplicate | Mother match |
| 1157 / 890 | 1157 | Deduplicate | Mother match |
| 1176 / 329 | 1176 | Deduplicate | Sex mismatch |
| 1218 / 1182 | 1218 | Deduplicate | Offspring match |
| 1276 / 1765 | 1765 | Deduplicate | Sex mismatch |
| 1542 / 7082 | 1542 | Deduplicate | Mother match |
| 1606 / 1499 | 1606 | Deduplicate | Mother match |
| 3017 / 3503 | 3017 | Deduplicate | Mother match |
| 3643 / 4882 | 3643 | Deduplicate | Sex mismatch |
| 3695 / 1271 | 3695 | Deduplicate | Mother match |
| 478 / 3444 | 3444 | Deduplicate | Sex mismatch |
| 4883 / 1607 | 1607 | Deduplicate | Sex mismatch |
| 4889/ 4933 | 4933 | Deduplicate | Mother match |
| 4961 / 5550 | 4961 | Deduplicate | Sex mismatch |
| 498 / 1303 | 498 | Deduplicate | Offspring match |
| 5542 / 6136 | 6136 | Deduplicate | Sex mismatch |
| 5546 / 6175 | 6175 | Deduplicate | Sex mismatch |
| 5563 / 5528 | 5528 | Deduplicate | Mother match |
| 5567 / 5529 | 5529 | Deduplicate | Mother match |
| 5585 / 5613 | 5613 | Deduplicate | Sex mismatch |
| 5586 / 6140 | 5586 | Deduplicate | Mother match |
| 5828 / 5624 | 5624 | Deduplicate | Mother match |
| 6145 / 5636 | 5636 | Deduplicate | Mother match |
| 6150 / 5786 | 5786 | Deduplicate | Sex mismatch |
| 6158 / 6142 | 6142 | Deduplicate | Mother match |
| 6165 / 5516 | 5516 | Deduplicate | Mother match |
| 6176 / 5551 | 5551 | Deduplicate | Mother match |
| 6862 / 6670 | 6670 | Deduplicate | Sex mismatch |
| 864 / 3871 | 864 | Deduplicate | Offspring match |

b) Sex-mismatched samples
| Sample ID | Candidate Sample ID | Action | Evidence |
| --- | --- | --- | --- |
| 774 | Unknown | Exclude | No pedigree evidence |
| 3088 | 5145 | Swap | Pedigree match |
| 5145 | 3088 | Swap | Pedigree match |
| 6156 | 6160 | Swap | Pedigree match |
| 6160 | 5585 | Swap | Pedigree match |
| 6166 | 5546 | Swap | Pedigree match |
| 1229 | Unknown | Exclude, re-sex sample | No pedigree evidence |
| 1435 | Unknown | Exclude | No pedigree evidence |
| 1676 | Unknown | Exclude, re-sex sample | No pedigree evidence |
| 5554 | 5511 | Swap | Pedigree match |
| 5555 | 5512 | Swap | Pedigree match |
| 5827 | 5831 | Swap | Pedigree match |
| 6049 | 6165 | Swap | Pedigree match |
| 6149 | 6147 | Swap | Pedigree match |
| 1668 | 1668 | Include, change sex | Pedigree match |
| 4893 | 4893 | Include, change sex | Pedigree match |
| 5512 | 6151 | Exclude | No pedigree evidence |
| 5517 | 5836 | Swap | Pedigree match |
| 5552 | 6151 | Exclude | No pedigree evidence |
| 6063 | Unknown | Exclude | No pedigree evidence |

c) Pedigree mismatched samples
| Sample ID | Candidate Sample ID | Action | Evidence |
| --- | --- | --- | --- |
| 5541 | 6149 | Swap | Pedigree match |
| 5562 | 5534 | Swap | Pedigree match |
| 5831 | 5563 | Swap | Pedigree match |
| 5832 | 5562 | Swap | Pedigree match |
| 6151 | 6178 | Swap | Pedigree match |
| 6160 | 5585 | Swap | Pedigree match |
| 6178 | 6176 or 6063 | Exclude | No pedigree evidence |
| 6147 | 5517 | Swap | Pedigree match |
| 5511 | 6150 | Swap | Pedigree match |
| 5534 | 6145 | Swap | Pedigree match |
| 5787 | Unknown | Exclude | No pedigree evidence |
| 5687 | 5538 | Swap | Pedigree match |
| 5836 | Unknown | Exclude | No pedigree evidence |
| 5781 | 6140 | Exclude | No pedigree evidence |
| 5538 | 6166 | Swap | Pedigree match |
| 890 | 890 | Keep | Pedigree match |
| 1273 | 1273 | Keep | Pedigree match |
| 1269 | 1269 | Keep | Pedigree match |

**Table S8.** Evaluating parentage assignment conflicts by MasterBayes (using microsatellites) and sequoia (using 572 SNPs) for 35 sample-verified individuals using recorded meta-data: territory, breed group, and genomic relatedness comparisons.

| Offspring ID | MasterBayes ID (dam, sire) | sequoia ID (dam, sire) | Dam match | Sire match | Consensus |
| --- | --- | --- | --- | --- | --- |
| 1202 | 469, 467 | 775, 467 | Mismatch | Match | Sequoia (relatedness) |
| 1230 | 469, 467 | 775, 467 | Mismatch | Match | Sequoia (relatedness) |
| 1268 | 931, 1040 | 914, 1040 | Mismatch | Match | Sequoia (breed group) |
| 1396 | 469, 467 | 775, 467 | Mismatch | Match | Sequoia (relatedness) |
| 1554 | 932, 1274 | 308, 1274 | Mismatch | Match | Sequoia (breed group) |
| 1761 | 1186, 921 | 604, 921 | Mismatch | Match | NA sequoia mum not seen from two years before, but possibly missed this bird during fieldwork? |
| 4944 | 3648, 1777 | 1872, 1777 | Mismatch | Match | Sequoia (breed group) |
| 4973 | 2968, 1553 | 1384, 1553 | Mismatch | Match | Sequoia (relatedness) |
| 5320 | 5263, 1795 | 1824, 1795 | Mismatch | Match | Sequoia (breed group) |
| 5602 | 1803, 1542 | 1580, 1542 | Mismatch | Match | Sequoia (relatedness, territory, caught as adult) |
| 5621 | 3616, 2966 | 1763, 2966 | Mismatch | Match | Sequoia (relatedness, territory, caught as adult) |
| 6031 | 5596, 5249 | 3596, 5249 | Mismatch | Match | Sequoia (relatedness, territory, caught as adult) |
| 6069 | 3518, 1869 | 5825, 1869 | Mismatch | Match | Sequoia (relatedness, territory, caught as adult) |
| 6742 | 6727, 5590 | 3462, 5590 | Mismatch | Match | Sequoia (territory, caught as adult) |
| 6773 | 6186, 4861 | 4861, 2974 | Mismatch | Match | Sequoia (relatedness, territory, caught as adult) |
| 884 | 469, 467 | 775, 467 | Mismatch | Match | Sequoia (relatedness, territory, caught as adult) |
| 1304 | 677, 1168 | 677, 550 | Match | Mismatch | Sequoia (relatedness, territory) |
| 1538 | 459, 607 | 459, 176 | Match | Mismatch | Sequoia (territory) |
| 1794 | 1382, 1677 | 1382, 847 | Match | Mismatch | Sequoia (territory) |
| 1888 | 2987, 1676 | 2987, 1695 | Match | Mismatch | Sequoia (territory) |
| 3008 | 1605, 2966 | 1605, 497 | Match | Mismatch | Sequoia (territory, age) |
| 3650 | 1781, 1512 | 1781, 1871 | Match | Mismatch | Sequoia (relatedness) |
| 5071 | 1532, 1804 | 1532, 911 | Match | Mismatch | Sequoia (territory, breedgroup, relatedness) |
| 6222 | 5530, 6026 | 5530, 3089 | Match | Mismatch | Sequoia (territory nearby, relatedness) |
| 1193 | 563, 851 | NA, 917 | NA | Mismatch | Sequoia (relatedness) |
| 1211 | NA, 547 | 843, 642 | NA | Mismatch | Sequoia (territory, caught as adult) |
| 1224 | 838, 215 | 502, NA | Mismatch | NA | Sequoia (territory, relatedness) |
| 1235 | NA, 1039 | NA, 216 | NA | Mismatch | NA not enough info |
| 1272 | 855, 854 | 601, NA | Mismatch | NA | NA not enough info |
| 1717 | NA, 1360 | 915, 1412 | NA | Mismatch | Sequoia (territory, relatedness, caught as adult) |
| 1744 | NA, 1577 | 308, 921 | NA | Mismatch | Sequoia (territory nearby, relatedness, caught as adult) |
| 4803 | 1042, 453 | 869, NA | Mismatch | NA | Sequoia (territory, relatedness, caught as adult) |
| 5849 | 5123, 1577 | NA, 5775 | NA | Mismatch | NA not enough info |
| 852 | NA, 559 | NA, 674 | NA | Mismatch | NA not enough info |

## Supplementary figures

**Figure S1.**
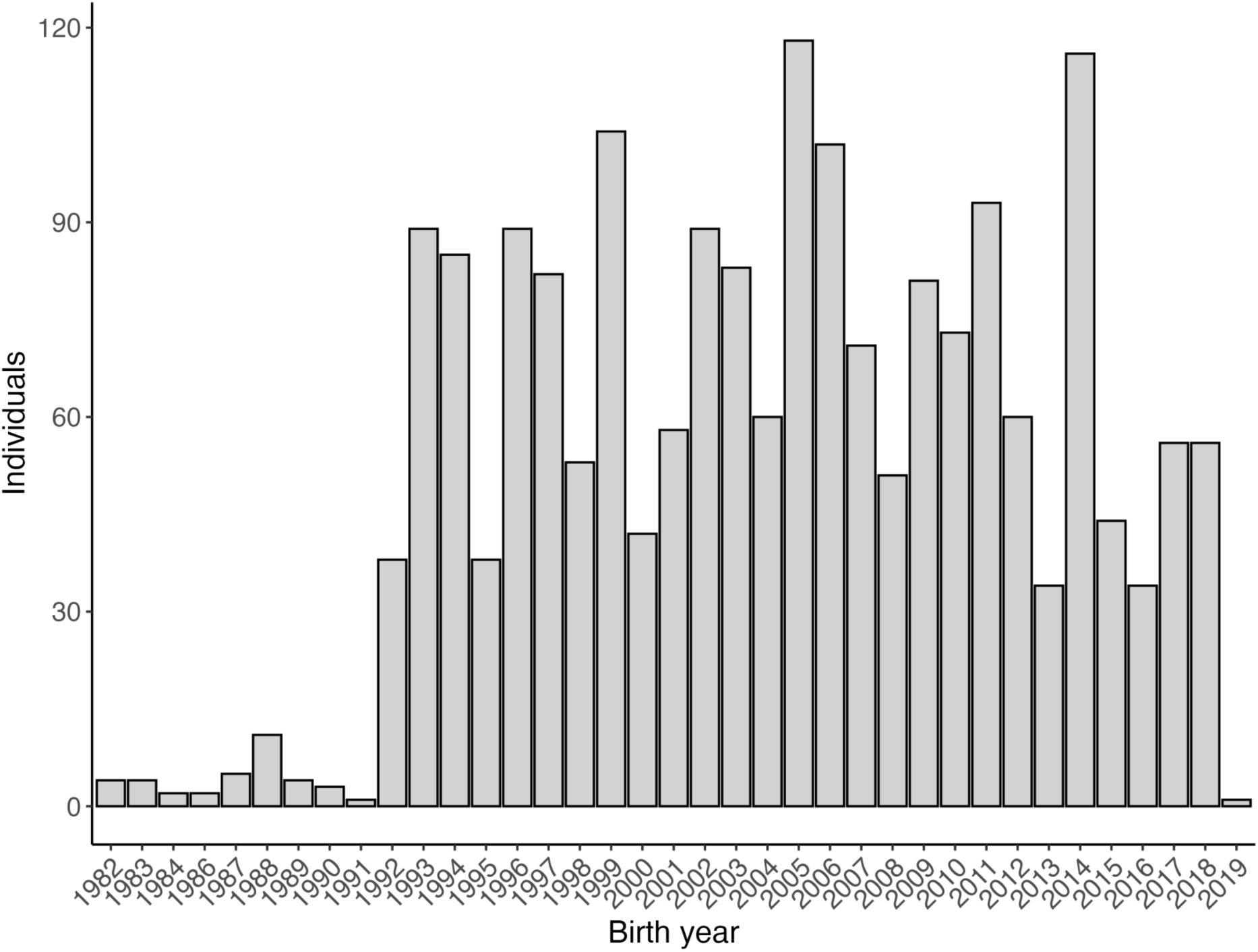
Numbers of individual Seychelles warblers whole-genome sequenced, by birth year, from the long-term sampled Cousin Island population.

**Figure S2.**
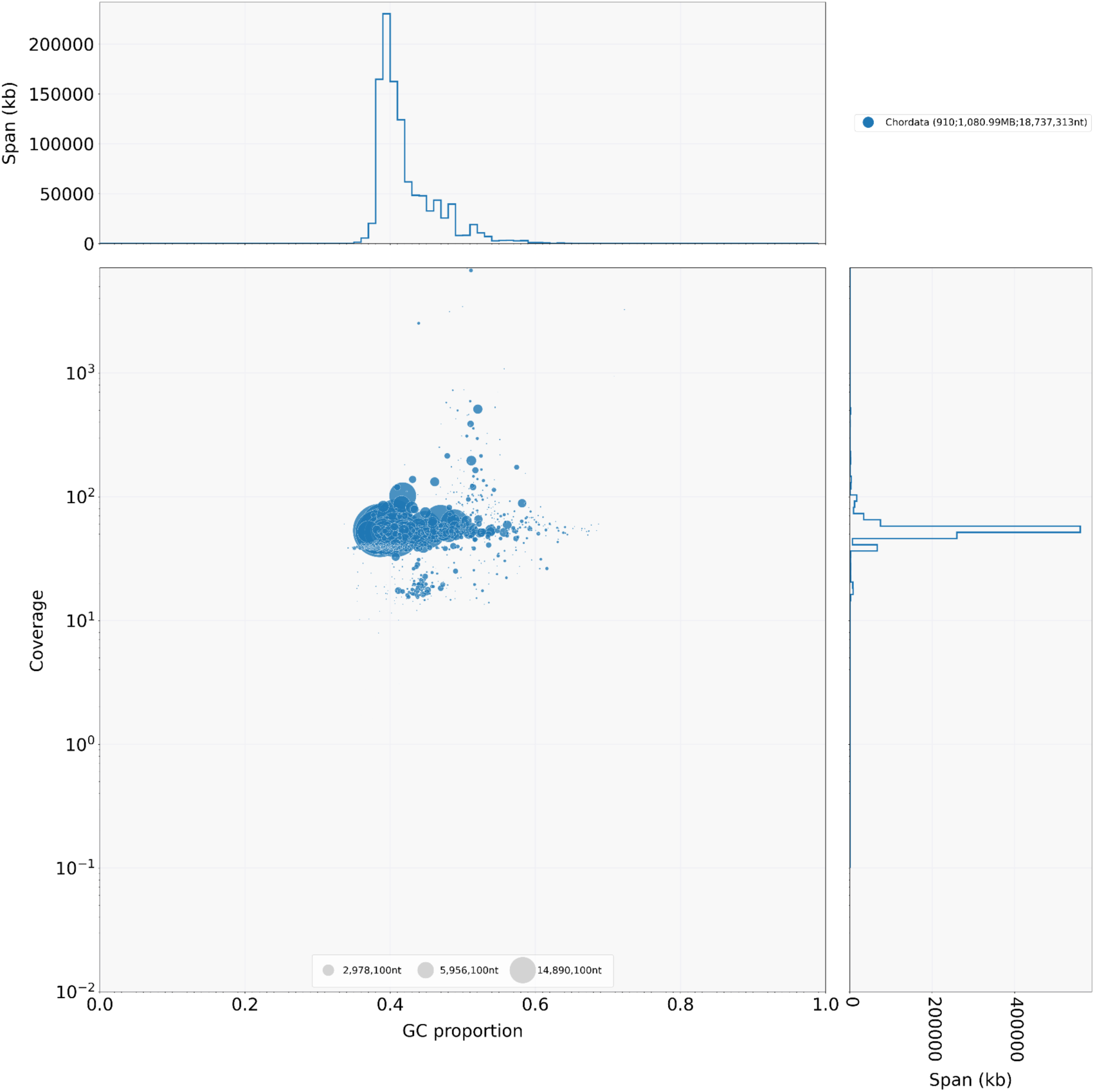
Blobtools contamination scan of Seychelles warbler genome assembly.

**Figure S3.**
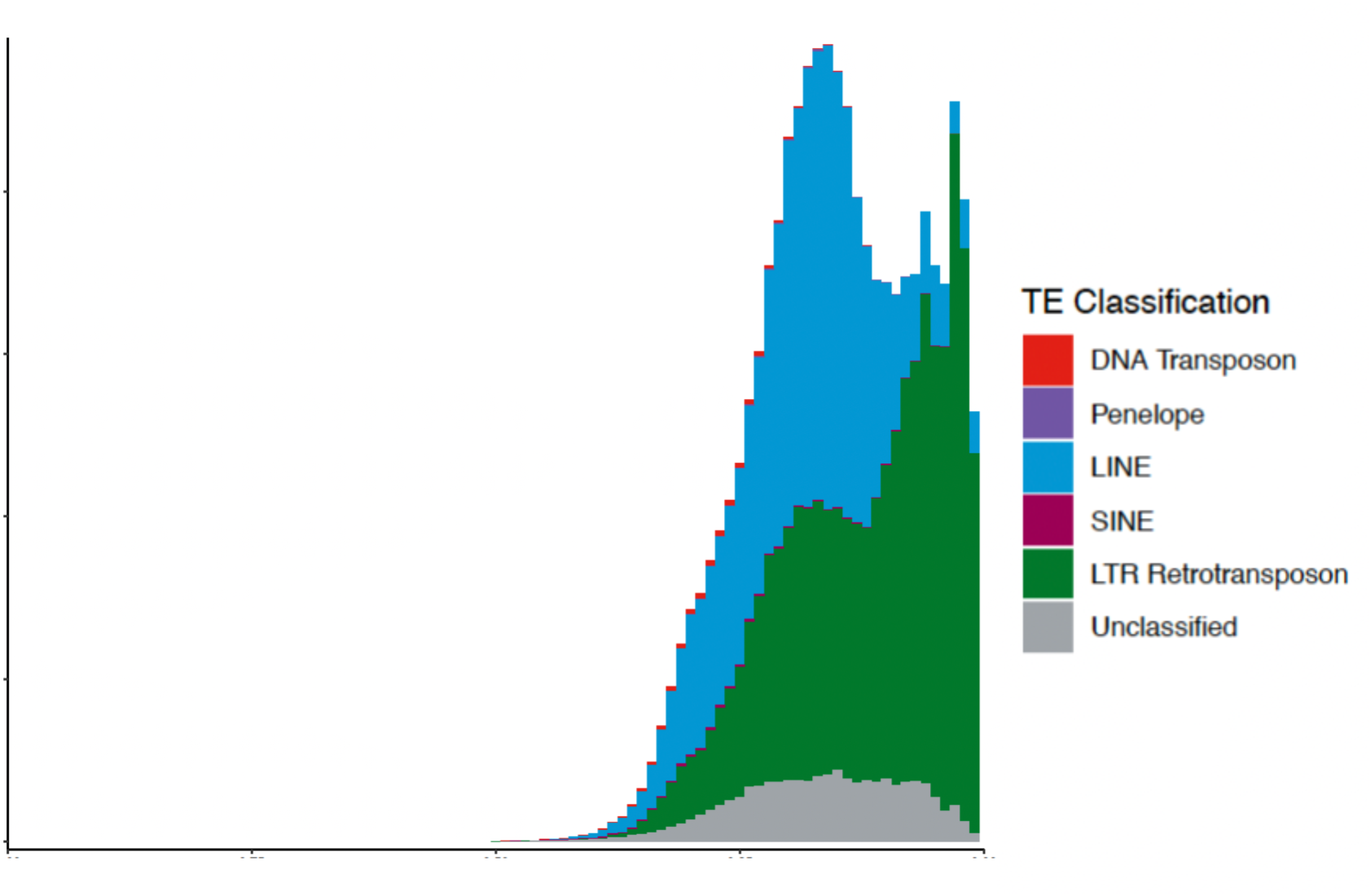
Repetitive DNA landscape in the female Seychelles warbler genome

**Figure S4.**
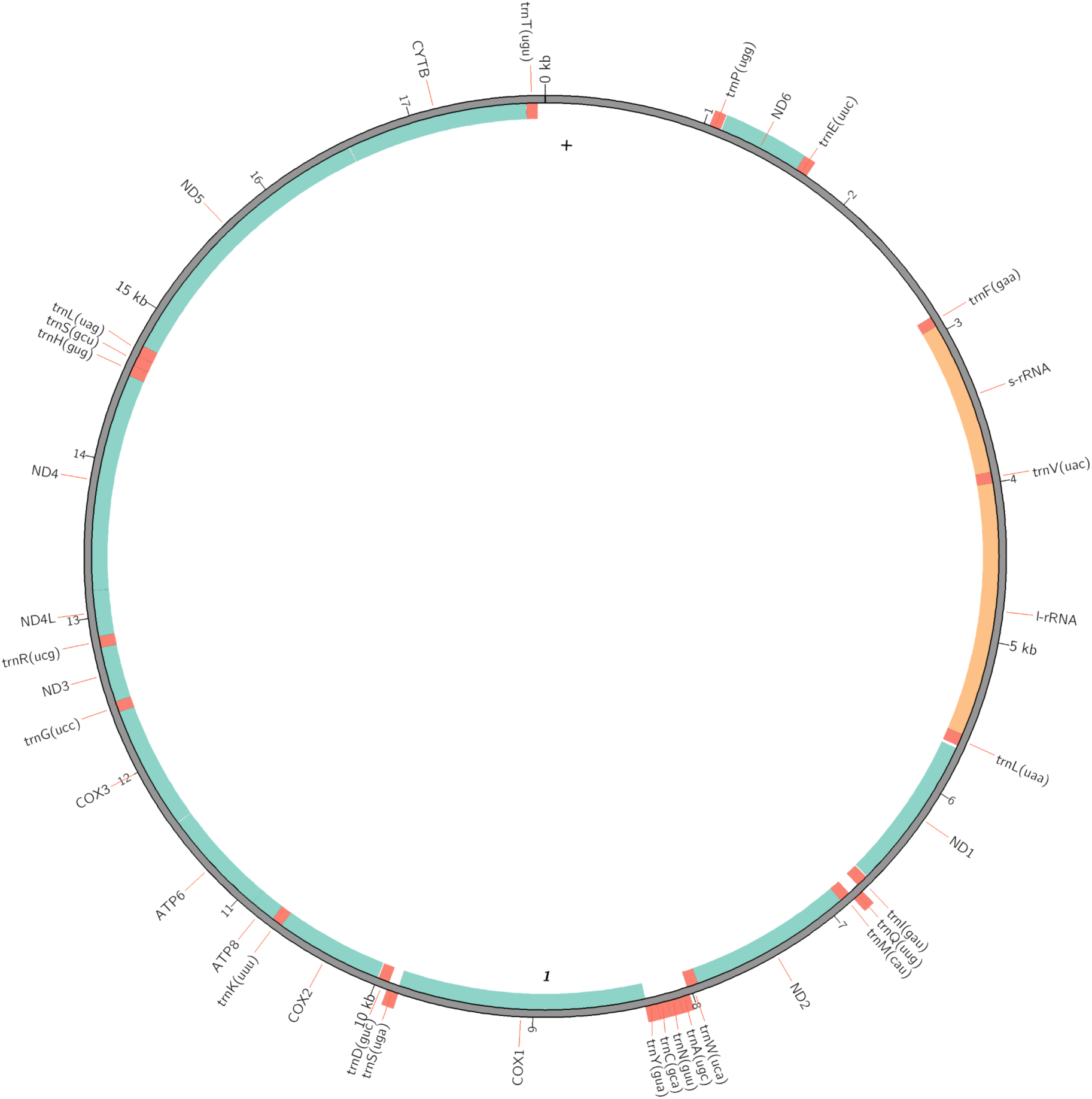
The complete, annotated, circularised mitochondrial genome is 17,847 bp, represented here as a Circos plot.

**Figure S5.**
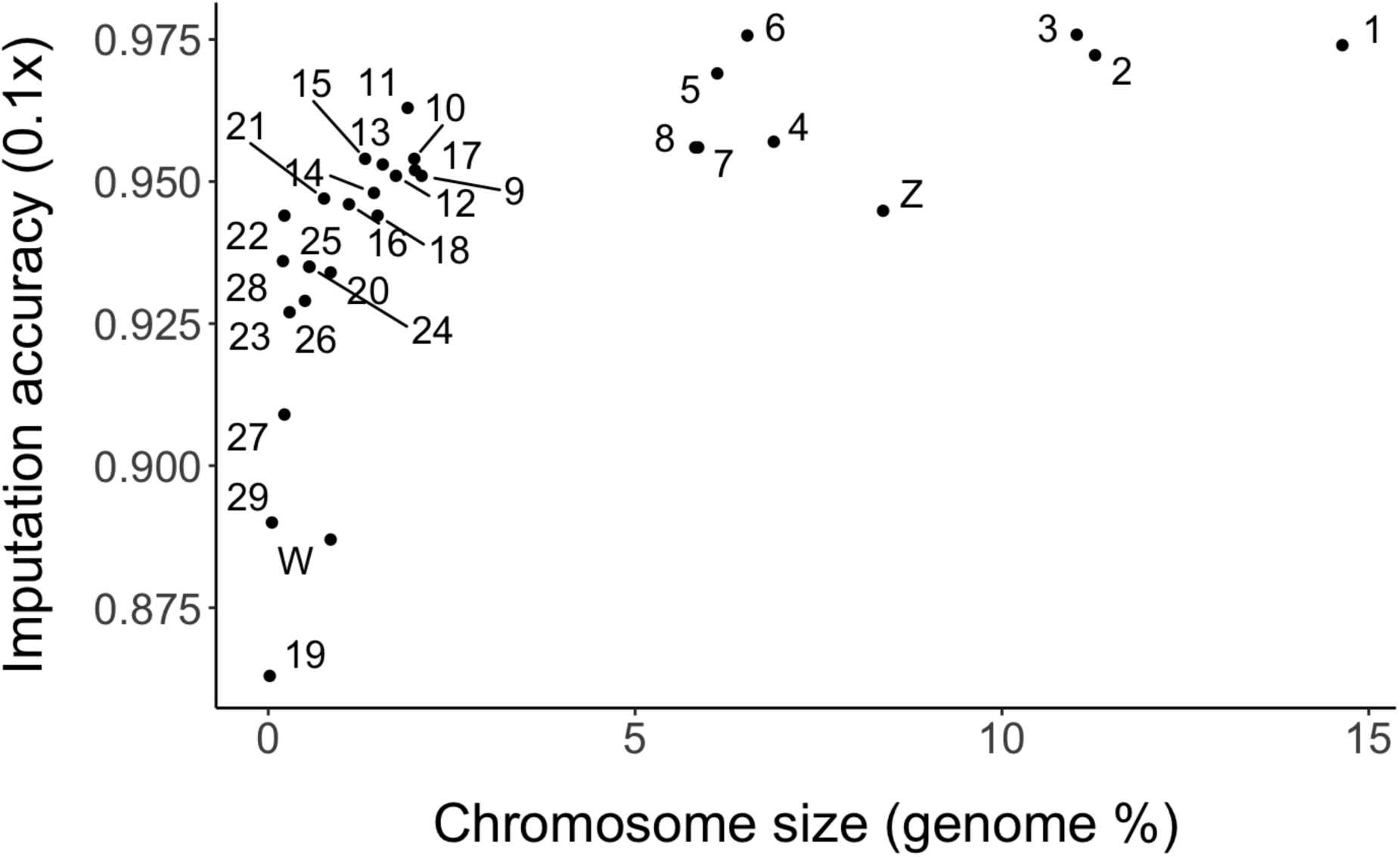
Imputation accuracy of STITCH per chromosome, estimated by comparing genotypes imputed from a downsampled 0.1× test set against the original high-coverage reference genotypes for the same individuals.

## Author contributions

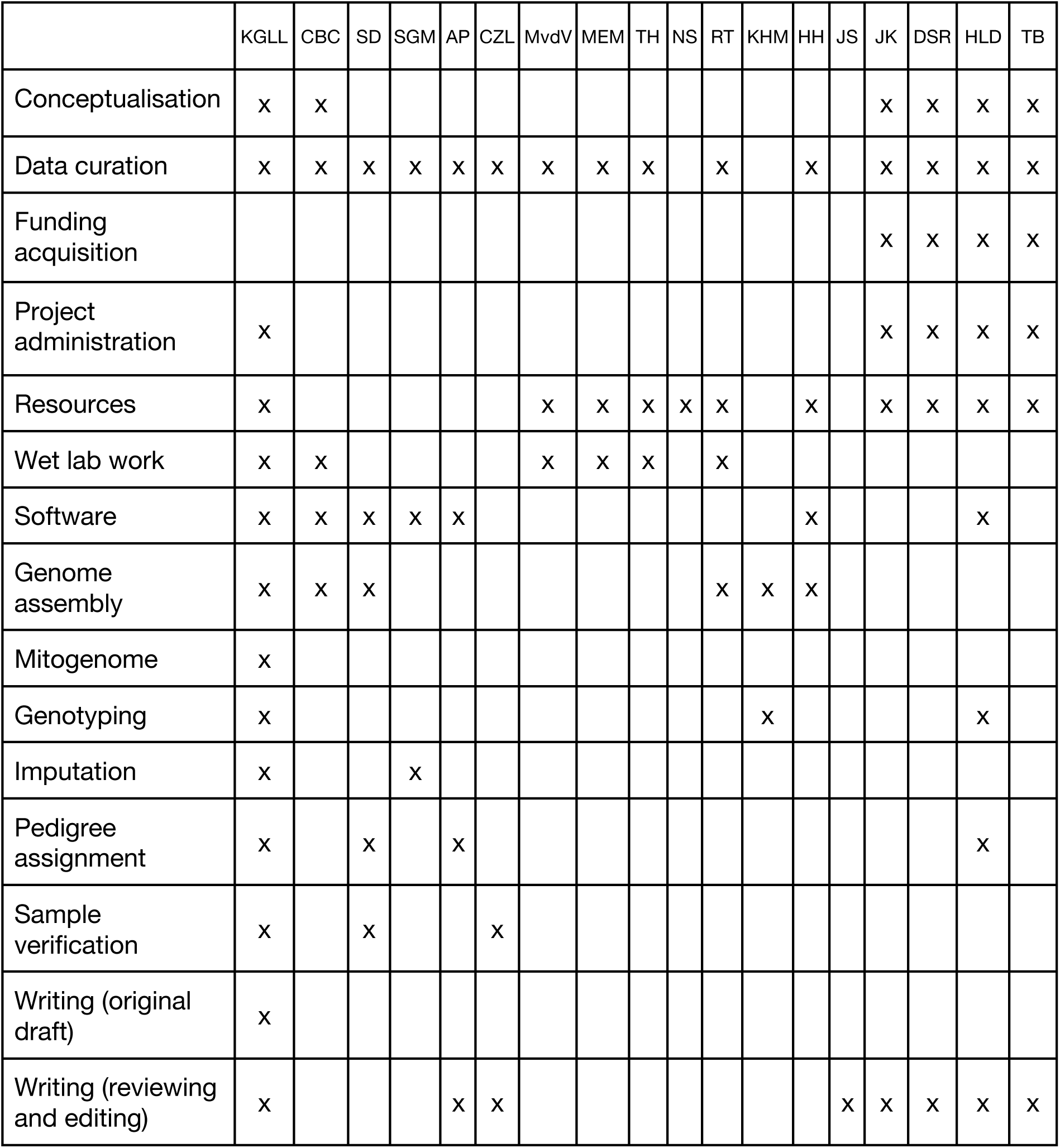

## Acknowledgements

We thank the following:

Nature Seychelles and their wardens for facilitating research on Cousin Island.

Seychelles Bureau of Standards and the Ministry of Agriculture, Climate Change and Environment for providing permission to conduct fieldwork and sample collection.

Fieldworkers, laboratory technicians, and database managers currently and previously associated with the Seychelles Warbler Project.

The IT Services at The University of Sheffield for the provision of services for High Performance Computing and troubleshooting and Hábrók at the University of Groningen.

NERC Environmental Omics Facility (NEOF) for sequencing and bioinformatics support. We appreciate, in particular, the support of Rowan Connell (University of Liverpool), Katy Maher, Ewan Harney, Gavin Gouws, Tom Holden, Lucy Knowles and Celine Pagnier (all at the University of Sheffield).

Natural Environment Research Council (NERC) and the Dutch Research Council (NWO) for the following grants, but also many past grants that have supported the long-term monitoring of the Seychelles warbler population:

NERC NE/P011284/1 to H.L.D. and D.S.R for genome sequencing.

NERC NE/B504106/1 to T.A.B. and D.S.R., NWO Rubicon 825.09.013 and NERC

NE/I021748/1 fellowships to H.L.D., and NWO visitors grant 040.11.232 to J.K. and

H.L.D. for previous pedigree work.

## Conflict of interest statement

The authors declare no conflict of interest.

## Data availability

Scripts can be found at: https://zenodo.org/records/18500278. The genome assemblies and individual whole-genome sequence data can be found temporarily at: https://zenodo.org/records/14717915 and permanently after one year’s embargo from date of publication at European Nucleotide Archive, under accession ID: PRJEB100611. We kindly suggest prospective researchers complete the following form before using the genomic data: https://docs.google.com/forms/d/e/1FAIpQLSfHHWT7XcyZtTNbI0wUYv3sw3XxI9G1AzhXB <u>SvlH-oyKYGrJA/viewform?usp=dialog</u>. There are a few reasons for this: 1) to help recognise local sovereignty of Seychelles warbler data, 2) because the data are subject to change, 3) someone may be performing your proposed analysis who can help.

